# Unraveling the mesoscale resting-state functional connectivity of ocular dominance columns in humans using high-resolution functional MRI

**DOI:** 10.1101/2025.03.27.645795

**Authors:** Marianna E. Schmidt, Iman Aganj, Jason Stockmann, Berkin Bilgic, Yulin Chang, W. Scott Hoge, Evgeniya Kirilina, Nikolaus Weiskopf, Shahin Nasr

**Author notes:** **Co-first authors** (Marianna E. Schmidt, Iman Aganj). **Corresponding author:** Shahin Nasr, Ph. D. **Email address:**.

## Abstract

Despite their importance for visual perception, functional connectivity between ocular dominance columns (ODCs)in the human primary visual cortex (V1) remains largely unknown. Using high-resolution functional MRI (fMRI), we localized ODCs and assessed their resting-state functional connectivity (rs-FC) in 12 human adults. Consistent with anatomical studies in animals, we found stronger rs-FC in the middle compared to deep and superficial cortical depths and selectively stronger rs- FC between ODCs with alike compared to unalike ocular polarity. Beyond what was known from animal models, and consistent with human perceptual biases, intra- and interhemispheric rs-FC was stronger in peripheral (compared to central) and in dorsal (compared to ventral) V1 subregions. Lastly, rs-FC patterns correlated with ODC maps, suggesting that ODC maps can be predicted from rs-FC patterns within V1. These results highlight the heterogeneity in rs-FC between ODCs across cortical depths and V1 subfields, underscoring their likely association with human perceptual features.

## 1. Introduction

In humans and many animals, the visual cortex is organized into neuronal columns that encode visual features. Despite their central role in shaping perception, the functional connectivity between cortical columns in the human visual cortex remains poorly understood. This gap in our knowledge stems primarily from the limited spatial resolution of conventional neuroimaging techniques relative to the size of cortical columns and layers. As a result, much of our understanding of the mesoscale organization of the visual cortex relies on anatomical studies in animals using invasive techniques that are not applicable to humans in vivo. To bridge this knowledge gap, we leveraged high-resolution functional MRI (fMRI) to investigate resting-state functional connectivity (rs-FC) between ocular dominance columns (ODCs), which are considered the building blocks of visual processing in the primary visual cortex (V1).

Anatomical studies in non-human primates (NHPs) have revealed that ODCs as far as 8 mm apart are systematically interconnected via horizontal fibers^1–6^. These fibers are most dense in the middle cortical layers, particularly layer 4B^6^, preferentially connecting ODCs with alike rather than unalike ocular polarity^4,7,8^. The level of this bias in the number of horizontal fibers remains a matter of debate varying from 4%^4^ to 15%^7^ in favor of the ODC pairs with alike ocular polarity. However, the reason for this inconsistency could not be tested systematically due to limited sample sizes.

Functional evidence for the biased structural connectivity between alike ODCs is provided by electrophysiological^9–11^ and optical imaging studies^12^. According to these studies, neurons with alike rather than unalike ocular polarity exhibit stronger synchronized activity. Interestingly, this synchronous activity is detectable over distances up to 20 mm^12^ indicating that the selective functional connectivity between ODCs extends far beyond the range of monosynaptic horizontal fibers and cannot be attributed to the overlapping receptive fields of the targeted neurons^11^.

To our knowledge, evidence for homologous functional organization within human V1 has not yet been reported. This is mainly due to difficulties in studying ODCs based on conventional neuroimaging techniques that provide a low signal-to-noise ratio and poor spatial resolution (>2mm), inadequate for studying the mesoscale functional organization of the human visual cortex. Nonetheless, at coarser levels, fMRI studies have shown higher intra- and interhemispheric rs-FC between V1 subregions representing similar locations of the visual field^13–15^. Outside V1, high-resolution fMRI has successfully been used to study rs-FC between color-, stereo- as well as motion-selective cortical columns across extrastriate visual areas^16–19^. Findings from these studies have supported the hypothesis that cortical columns with alike stimulus preferences are more selectively connected to each other, and that this selectivity can be detected by high-resolution resting-state fMRI (see also^20^).

Using high-resolution fMRI in the past, multiple groups have successfully localized ODCs in humans^16,21–27^. In this study, we extended the application of high-resolution fMRI to study the rs-FC between ODCs, in the absence of any visual input (eyes closed), in 12 adult humans. We examined the influence of cortical depth, pairwise distance, relative ocular polarity (i.e., alike vs. unalike ocular preference), and strength of ocular preference on the mesoscale rs-FC within V1 and assessed the heterogeneity of the measurements across V1 subregions. These analyses were then extended to evaluate the mesoscale rs-FC between V1 subregions within and between the two hemispheres. Lastly, we tested the predictability of ODC maps based on the mesoscale rs-FC pattern within V1.

In agreement with animal studies, we found selective mesoscale rs-FC between ODCs with stronger connectivity at middle compared to deep and superficial cortical depth levels. Beyond what was previously known from animal studies, we showed that these connectivity patterns are not uniform but vary across V1 subregions within and between hemispheres, consistent with the heterogeneity of visual processing across subfields. Finally, we demonstrated that rs-FC patterns correlate with ODC maps, providing an initial step toward predicting and segmenting the mesoscale functional organization of the human visual cortex. A preliminary version of the results has been already presented at a workshop^28^.

## 2. Results

Studies in NHPs suggest that horizontal fibers in primate V1 are more numerous in the middle layer (layer 4B) compared to deeper (layers 5 and 6) and superficial cortical layers (layers 2 and 3)^6^. They also indicate a significant difference (4% to 15%) in the number of fibers connecting ODCs, favoring pairs with alike rather than unalike ocular polarity^4,7,8^. Here, we first examined whether these mesoscale features are also reflected in rs-FC between vertex pairs in human V1. Then, we extended our analysis to investigate the influence of ocular preference strength, the heterogeneity of these features across different V1 subregions and their presence in interhemispheric rs-FC measurements.

To achieve these aims, we acquired high-resolution fMRI data at 1 mm isotropic resolution in the visual cortex of 12 adults (Figure 1, see Methods). ODCs were localized using dichoptic stimulation and resting-state fMRI data were collected in a separate session. Individual visual area borders and central, peripheral, dorsal, and ventral V1 subregions were defined using retinotopic mapping (eccentricity <10°), and rs-FC was quantified by correlating resting-state time series between cortical vertices within deep, middle and superficial cortical depth levels of V1. Connectivity was then assessed as a function of pairwise distance, ocular polarity (alike vs. unalike), cortical depth, pairwise ocular preference strength, visual field representation (central vs. peripheral, dorsal vs. ventral), and interhemispheric connections. Finally, rs-FC patterns were correlated with ODC maps to test their correspondence.

**Figure 1.**
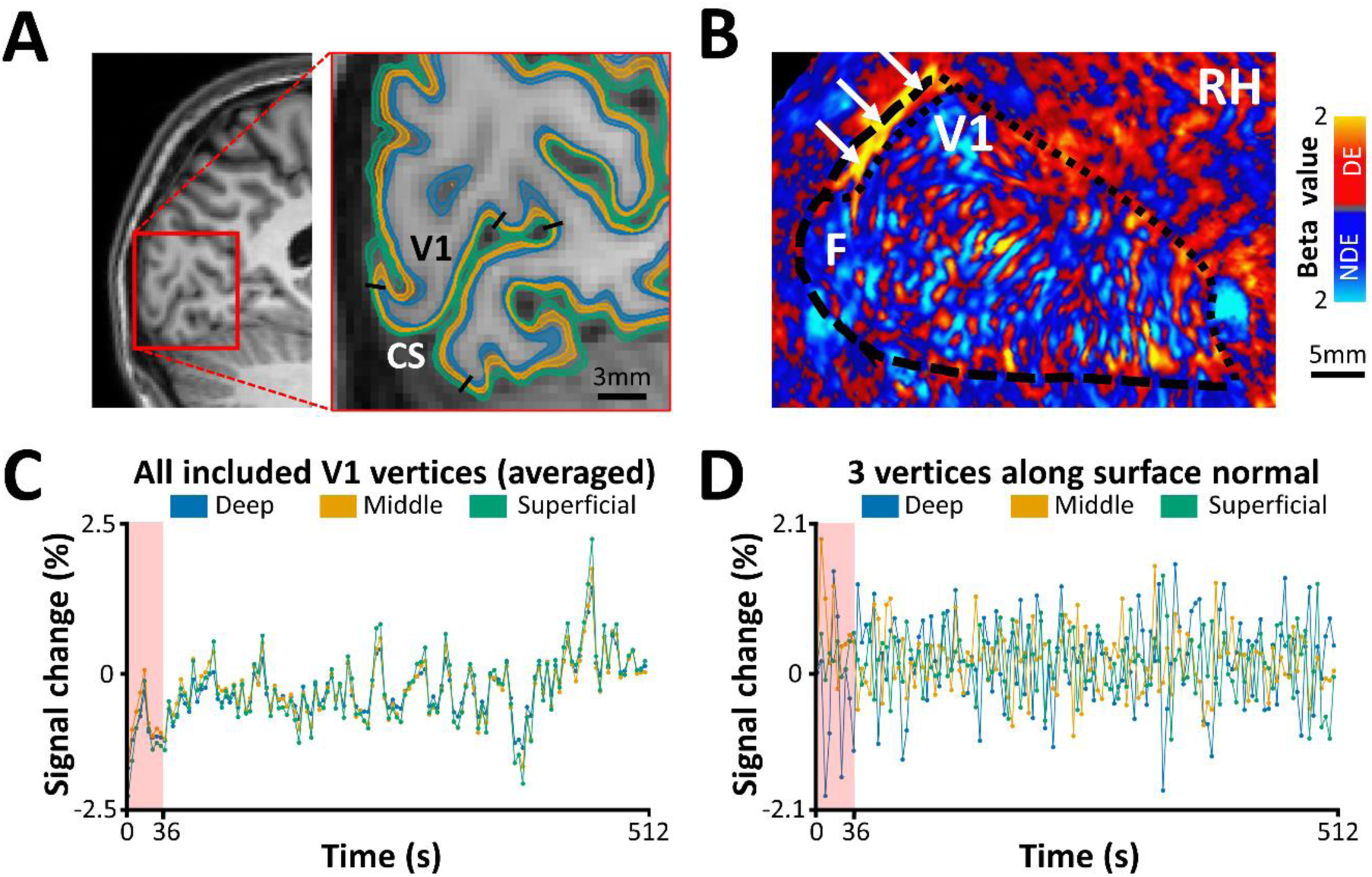
Structural and functional MRI from one individual participant. **A)** Structural scan and reconstruction of cortical surfaces at different depth levels. For each participant, primary visual cortex (V1) was localized retinotopically along the calcarine sulcus (CS). Black lines indicate the borders of the stimulated portion of V1 (radius<10°). As a part of brain reconstruction, the cortical gray matter was divided into deep (blue), middle (orange), and superficial (green) depths with small gaps between them (see Methods). **B)** Differential ocular dominance column (ODC) map (beta values) overlaid on the flattened cortex. The ocular dominance response was derived by contrasting the response to stimulation of the dominant eye (DE; red to yellow) versus the non-dominant eye (NDE; blue to cyan). The map shows the stimulated portion of V1 (radius<10°) in the right hemisphere (RH) of a representative subject. Notably, a large activity patch along the dorsal part of the vertical meridian (white arrows), in which the ODCs could not be mapped due to technical limitations, was excluded from the analysis^23^. **C)** Mean spontaneous fMRI time course measured over the whole stimulated portion of V1. Consistent with previous studies, the signal intensity was weaker in deeper compared to more superficial depths. To exclude the potential signal instabilities at the beginning of the scan, the first 36 s were excluded from further analysis. **D)** Spontaneous fMRI time courses from three vertices sampled at deep, middle, and superficial layers, respectively, at a randomly selected single tangential location along the surface normal. Compared to the averaged signal, these individual laminar time courses exhibit lower temporal correlation, underscoring the likely presence of depth-specific fluctuations.

### 2.1. Rs-FC between alike and unalike ODCs

Figure 2A illustrates the rs-FC levels, measured during the eyes-closed condition (see Methods), for vertex pairs with alike and unalike ocular polarity as a function of distance between the two vertices of the pair. A two-way RM-ANOVA (distance and relative ocular polarity), applied to the measured rs-FC levels, revealed that rs-FC was significantly stronger between alike than unalike vertex pairs (F(1, 11)=6.12, *p*=0.03). Here while rs-FC decreased significantly with increasing distance between vertex pair (F(9, 99)=29.94, *p*<10^-3^), the interaction between the effects of distance and relative ocular polarity remained non-significant (F(9, 99)=0.58, *p*=0.58). These results suggest that selective rs-FC between vertices with alike ocular polarity persists across relatively long distances (Supplementary Figure 1)

**Figure 2.**
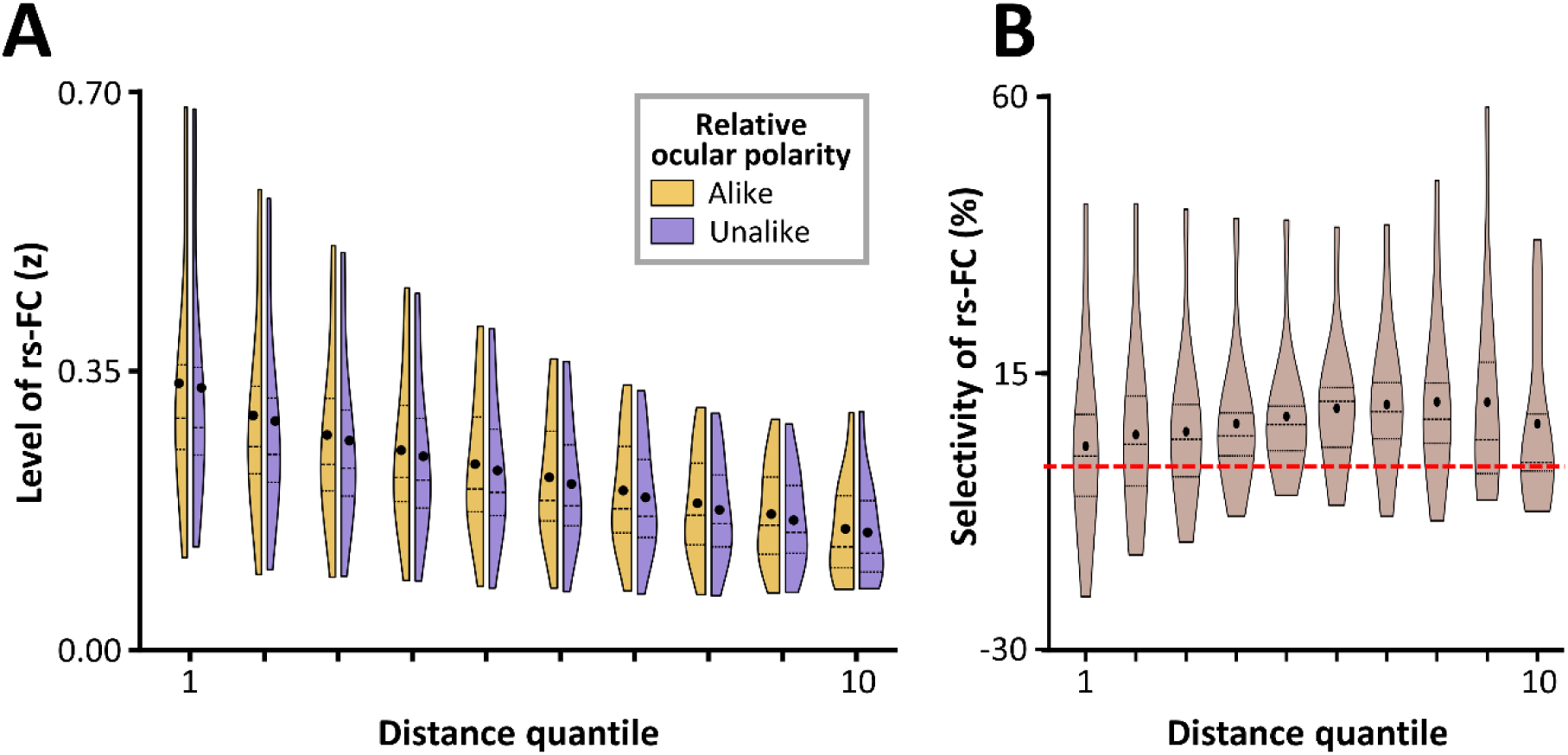
The resting-state functional connectivity (rs-FC) between vertex pairs in the primary visual cortex and its selectivity in favor of the pair with alike ocular polarity as a function of distance. **A)** The distribution of rs-FC (*z*-values) across subjects is displayed using split violin plots for vertex pairs with alike (orange) and unalike (violet) ocular polarity for 10 distance quantiles (with 1 being the closest but excluding vertices closer together than 3mm due to fMRI point spread function (see Methods) and 10 being the farthest distance (Supplementary Figure 1)). Rs-FC decreased with increasing distance. **B)** Selectivity of rs-FC, measured as the alike rs-FC minus the unalike rs-FC, is depicted similarly as violin plot for each distance quantile in percentage. To facilitate comparison with the previous findings in animal studies, values were normalized for overall rs-FC level. The red dashed line indicates 0. In both panels, the violins display the full distribution of the data, with thickness representing density and endpoints corresponding to the min and max outliers.

The absence of interaction between the effects of distance and relative ocular polarity is also apparent in Figure 2B that illustrates the level of rs-FC selectivity, measured as rs-FC between alike – unalike vertex pairs, in each distance quantile. To make the rs-FC selectivity measurement similar to those in the previous anatomical studies^4,7,8^, we normalized it relative to the overall level of rs-FC in the same quantile. As demonstrated here, rs-FC selectivity remained on average positive even at the highest tested distance quantile, where vertex pairs were approximately 35 mm apart (Supplementary Figure 1), far exceeding the expected reach of a single horizontal fiber^1,4,5^.

Notably, consistent with our previous reports in a smaller group of participants^23^, there was no significant difference between the number of vertices showing a preference for stimulation of the DE versus NDE (*p*=0.10) or their ocular preference level as measure by an ocular dominance index (ODI; see Methods) (*p*=0.53) (Supplementary Figure 2). Furthermore, a subsampling method assured us that the distance distribution was the same for alike and unalike vertex pairs (see Section 5.5.1.). Thus, these findings cannot be attributed to an unbalanced distribution of ocular dominance activity in V1.

### 2.2. Reproducibility of rs-FC measurements across scan sessions

We tested whether the selective rs-FC between alike ODCs could be reliably detected across different days. Four individuals were re-scanned on a separate day using a different ultra-high-field MRI scanner and a modified data acquisition protocol (see Section 5.3.2.). Here again, a three-way RM-ANOVA (distance, relative ocular polarity, and scan session) applied to the measured rs-FC across different distance quantiles (Supplementary Figure 3) confirmed robust effects of relative ocular polarity (F(1, 3)=25.46, *p*=0.02), and distance (F(9, 27)=19.62, *p*=0.02), while revealing no significant main effect of scan session (F(1, 3)=0.69, *p*=0.47) and no interactions between the effects of scan session and the other two factors (*p*>0.18). These results indicate that selective rs-FC between ODCs is reproducible over time and insensitive to scanner hardware and the exact acquisition parameters. Notably, the robustness of the ODC maps for our subjects has been tested and scrutinized in our previous studies^16,23^.

### 2.3. Selectivity of rs-FC as a function of ODI and cortical depth

Studies in NHPs have suggested that the strength of ocular preference as measured by ODI may influence the selectivity of horizontal fibers^4,7,8^. Accordingly, here, we tested the hypothesis that rs-FC depends not only on the relative ocular polarity of the vertex pair but also on their ODI. Furthermore, we tested whether the level of rs-FC varied across cortical depth levels, as suggested by animal models^6^, and if rs- FC selectivity was detectable across all cortical depths. Considering the absence of significant interaction between the effects of relative ocular polarity and the distance between vertex pairs, data from all distance levels were averaged in all further analyses.

Figure 3A illustrates the rs-FC between vertex pairs as a function of their ODI, relative ocular polarity, and cortical depth. A three-way RM-ANOVA revealed that rs-FC was significantly stronger between vertices with stronger ODIs than those with weaker ODIs (F(4, 44)=11.40, *p*<0.01) (Supplementary Table 1). As expected from our previous analysis, rs-FC was significantly stronger between vertex pairs with alike rather than unalike ocular polarity (F(1,11)=6.09, *p*=0.03). Here, a significant interaction between the effects of ODI and relative ocular polarity (F(4, 44)=6.08, *p*=0.02) demonstrated that rs-FC selectivity increases with the ODI of vertex pairs, reinforcing the link between functional connectivity and ocular dominance. This interaction is also apparent in Figure 3B that illustrates the level of rs-FC selectivity, measured as rs-FC between alike – unalike vertex pairs, without any normalization.

**Figure 3.**
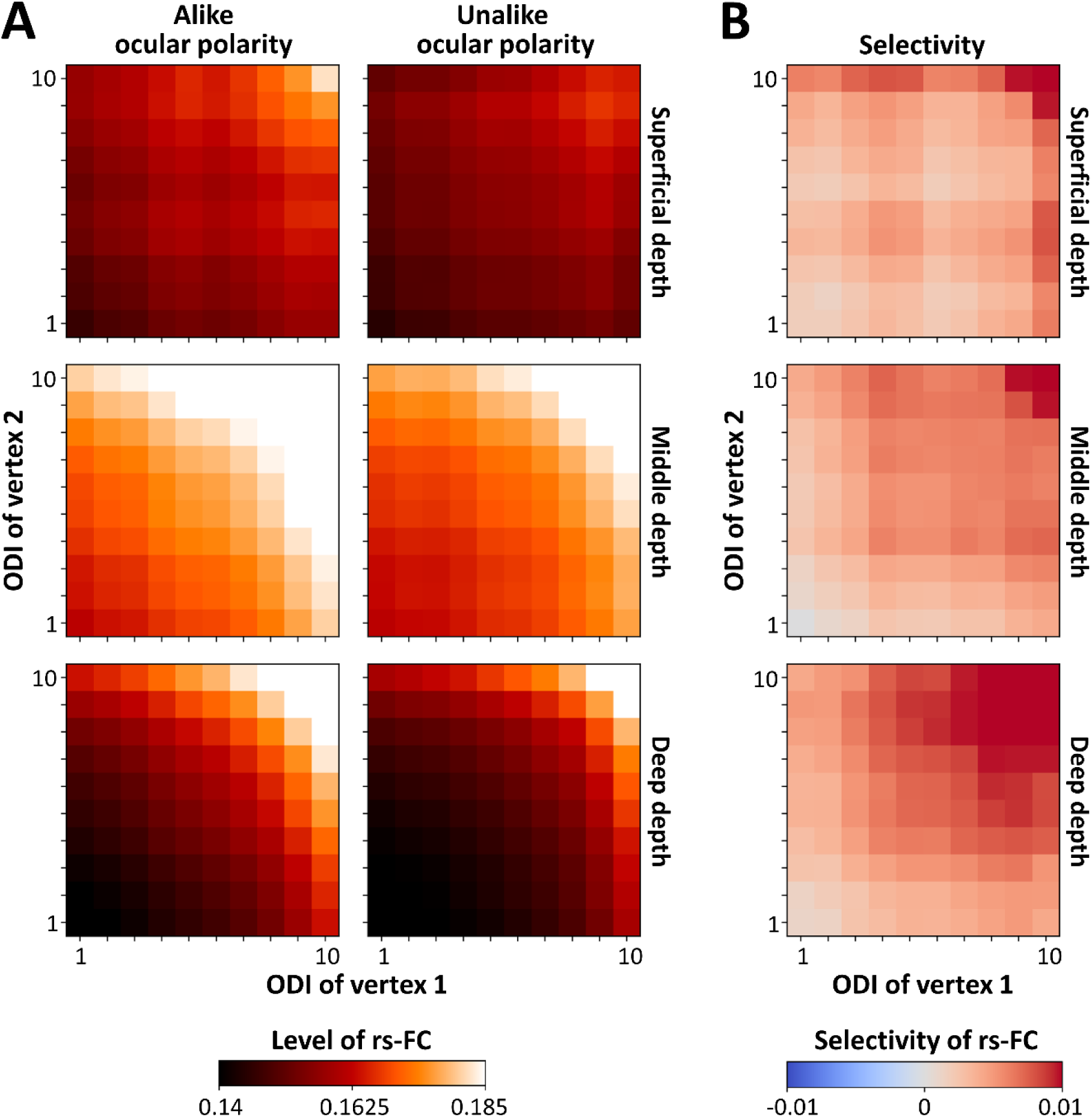
The measured resting-state functional connectivity (rs-FC) between primary visual cortex vertex pairs and its selectivity as functions of cortical depth and ocular dominance index (ODI) averaged over all participants. **A)** The level of rs-FC between vertex pairs with alike (left) and unalike (right) ocular polarity measured across superficial (top), middle (middle), and deep (bottom) cortical depths. Each correlation matrix shows the mean rs-FC measured as *z*-values. In each matrix, *x*- and *y*-axes represent the ODIs of vertex pairs categorized in quantiles 1 to 10 representing low to high ODI beta values (Supplementary Figure 6). The measured rs-FC is generally higher at middle cortical depth and increases with ODI. **B)** Selectivity is defined as the difference between rs-FC of vertex pairs (alike minus unalike ocular polarity) without any normalization. The selectivity was generally higher, favoring pairs with alike ocular polarity and increased for vertex pairs with higher ODI.

Moreover, consistent with the laminar organization of horizontal fibers in animals, rs-FC was highest in middle cortical depth compared to deeper and more superficial depths (F(2,22)=4.74, *p*=0.047). This effect was particularly pronounced in regions with stronger ODIs, resulting in a significant interaction between the effects of ODI and cortical depth (F(8, 88)=13.47, *p*<10^-4^). The lack of significant interaction between the effects of relative ocular polarity and cortical depth (*p*=0.30) suggests that rs-FC selectivity remains equivalent across cortical depths, thus ruling out the possibility that rs-FC selectivity was arbitrarily stronger simply because the ODC map was always sampled from deep cortical depth (to minimize the blurring effect of pial veins). Similar results were also found after applying an LME model to the rs-FC data (see Supplementary Analysis, and Supplementary Table 2).

Together, these results suggest that the influence of relative ocular polarity and cortical depth is contingent on higher ODIs. Considering the stronger fMRI signal in more superficial compared to deeper cortical depths across V1 (F(2,22)=74.23, *p*<10^-5^) (Figure 1C) and even within dorsal, ventral, central, and peripheral subregions individually (F(2,22)>49.98; *p*<10^-4^), these results also suggest that fMRI signal intensity does not dominate the strength of rs-FC measured at different cortical depths. Instead, other factors, such as the organization of horizontal fibers across cortical depths, may also influence the strength of rs-FC (see Discussion).

### 2.4. Heterogeneity in rs-FC

Multiple previous studies suggested stronger global visual processing and greater sensitivity to low spatial frequency components —essential for global visual processing—in the lower compared to the upper visual field^29–32^. Given that sensitivity to lower spatial frequency components requires sampling from a larger visual field, it could be argued that rs-FC may be stronger in dorsal V1 subregions representing the lower visual field compared to ventral V1 subregions representing the upper visual field. To test this hypothesis, we divided V1 into two equivalent ROIs, representing the upper and lower portions of the visual field (Figure 4A), and compared the rs-FC pattern across them.

**Figure 4.**
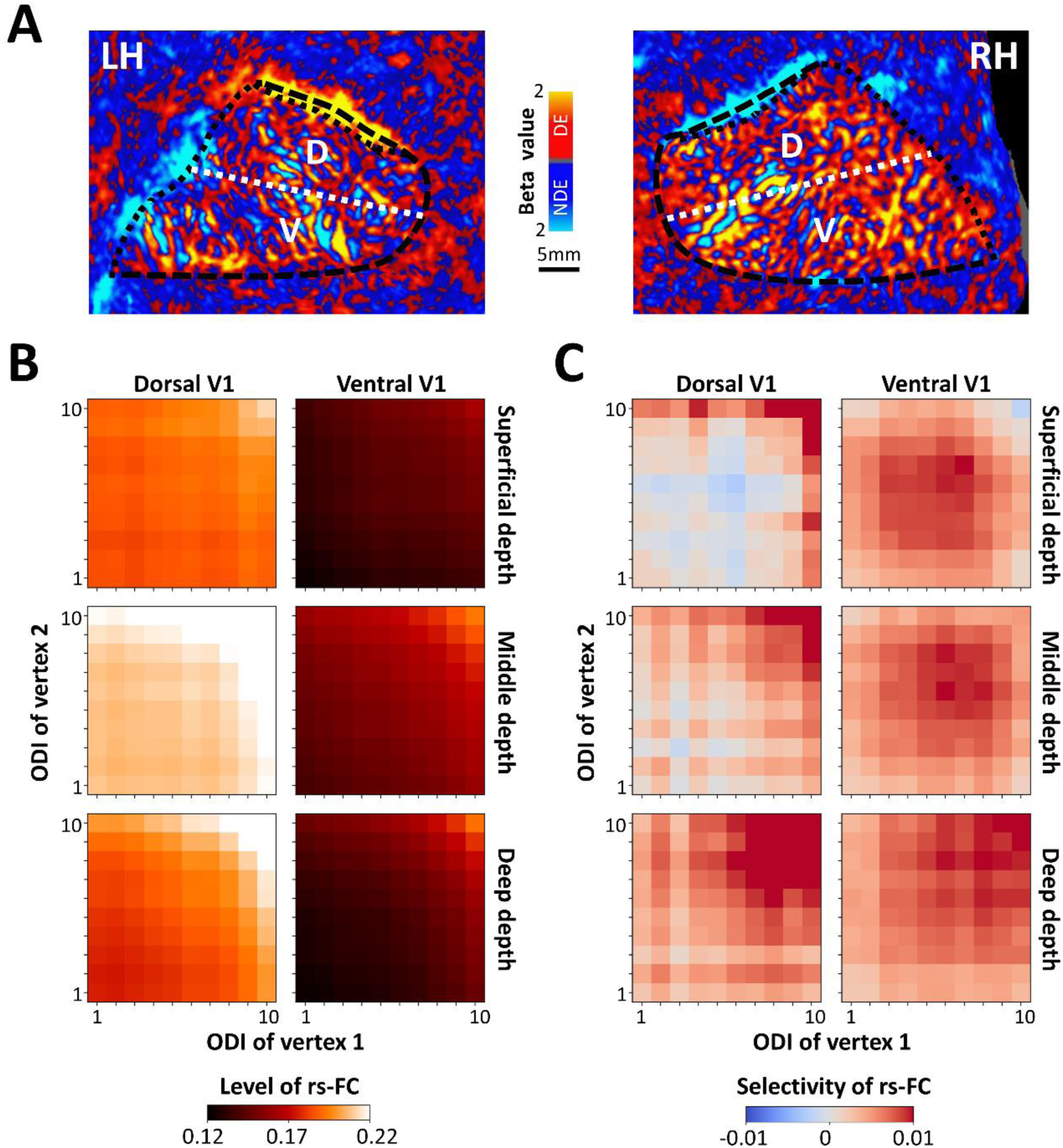
Comparison of resting-state functional connectivity (rs-FC) and selectivity between ventral (V) and dorsal (D) primary visual cortex (V1) averaged across all participants. **A)** Location of dorsal and ventral V1 subregions, overlaid on the ocular dominance column (ODC) map in the left and right hemispheres (LH and RH, respectively) from one participant (other than the one shown in Figure 1). The stimulated part of V1 (dashed and dotted black lines) was divided into two equally sized areas along the calcarine sulcus (dashed white line). **B)** The measured rs-FC between (all) vertex pairs in dorsal (left) and ventral (right) V1 subregions irrespective of relative ocular polarity. The rs-FC level was on average higher in dorsal compared to ventral V1 subregion. **C)** The selectivity of rs-FC across the two subregions. In dorsal V1 subregion, the selectivity was more confined to sites with high ocular dominance index (ODI) whereas in the ventral V1 subregion, the selectivity was more uniformly distributed across ODI. The other details are the same as in Figure 3.

Figure 4B illustrates the level of rs-FC measured in dorsal V1 (representing the lower visual field) and ventral V1 (representing the upper visual field). Overall, consistent with our hypothesis, rs-FC was stronger in the dorsal compared to ventral V1. Additionally, rs-FC selectivity was more uniform — i.e., independent of ODIs — in ventral compared to dorsal subregions (Figure 4C). A four-way RM-ANOVA (ROI, relative ocular polarity, ODI, and cortical depth) (Supplementary Table 3) applied to the measured rs-FC revealed a significant main effect of ROI (F(1, 11)=29.26, *p*<10^-3^), as well as significant interactions between the effects of ROI, relative ocular polarity, and cortical depth (F(2, 22)=5.17, *p*=0.03) as well as between the effects of ROI, ODI, and cortical depth (F(8, 88)=4.38, *p*=0.02). These findings indicate significant heterogeneity in both the level of rs-FC and its selectivity across dorsal and ventral V1 subregions, as expected from the differential visual processing mechanism in these regions.

A stronger preference for lower spatial frequency can also be found in V1 subregions representing peripheral compared to central visual fields^33–35^. Accordingly, we further tested if a similar heterogeneity can also be found between central and peripheral V1 subregions (Figure 5A). As demonstrated in Figure 5B, rs-FC was stronger in the peripheral compared to central V1 subregions across the superficial and middle (but not deep) cortical depths. The same pattern, but less pronounced, was found for the selectivity of rs-FC (Figure 5C). Here, the application of four-way RM-ANOVA (Supplementary Table 4) to the measured rs-FC showed a significant main effect of relative ocular polarity (F(1,11)=5.52; *p*=0.04) and a significant interaction between the effects of ROI and cortical depth (F(4, 44)=27.72, *p*=10^-4^). However, the two- and three-way interactions between the effects of ROI and relative ocular polarity remained non-significant (F<0.63, *p*>0.53). These results suggested that the overall pattern of rs-FC, its dependency on ODI, relative ocular polarity, and cortical depth may vary between V1 subregions, and that this heterogeneity appears to be consistent with the known biases in human visual perception across visual fields.

**Figure 5.**
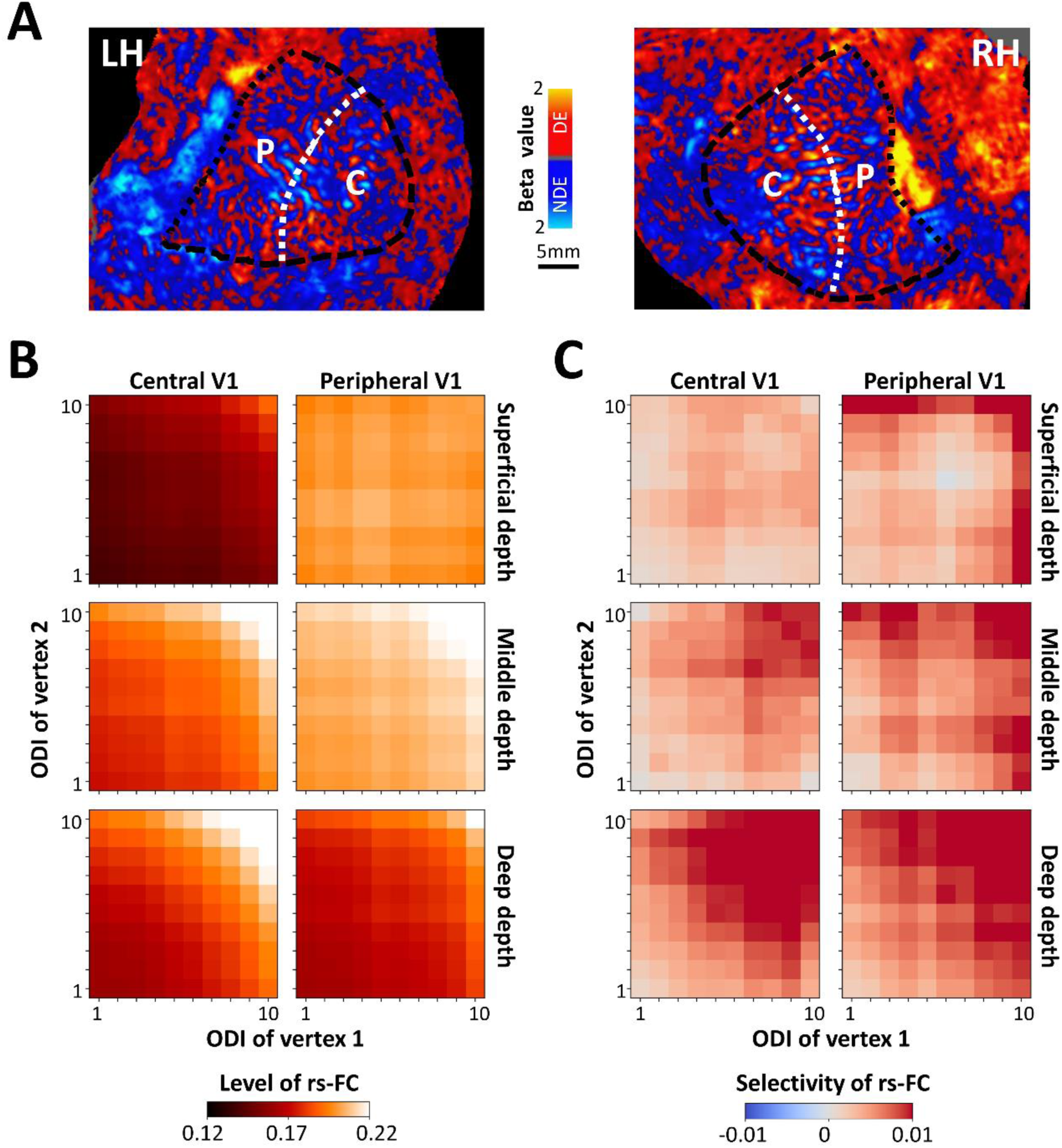
Comparison of resting-state functional connectivity (rs-FC) and selectivity between central (C) and peripheral (P) primary visual cortex (V1) averaged across all participants. **A)** Central (radius<4°) and peripheral V1 subregions are shown relative to the ocular dominance column (ODC) map in one participant (other than the one shown in Figures 1 and 4). **B)** The level of rs-FC was on average higher in peripheral V1 compared to central V1, which was more apparent in superficial and middle depths. **C)** Likewise, the selectivity of rs-FC was higher in the peripheral compared to the central V1 subregion in superficial and middle cortical depths. The other details are similar to Figure 4.

### 2.5. Interhemispheric rs-FC between ODCs

While monosynaptic callosal fibers directly linking V1 between hemispheres are limited^36–38^, numerous polysynaptic connections — primarily through feedback pathways from other callosally connected visual areas (e.g., V2) — may contribute to interhemispheric rs-FC in V1. Consistently, multiple studies have already shown interhemispheric rs-FC between V1 subregions that represent the same portion of visual field^13–15^. Here, we extended those findings by examining the dependence of interhemispheric rs-FC connecting V1 subregions (dorsal vs. ventral and central vs. peripheral) between the two hemispheres, on relative ocular polarity, ODI, and cortical depth. Because of the known retinotopic organization of interhemispheric connectivity, we limited our test to retinotopically corresponding areas, e.g., dorsal to dorsal and ventral to ventral V1.

As illustrated in Figure 6A, interhemispheric rs-FC exhibited at least two regional differences between dorsal and ventral V1. First, similar to intrahemispheric rs-FC, dorsal V1 showed stronger interhemispheric rs-FC than ventral V1. Second, rs-FC selectivity (Figure 6B) followed distinct patterns in these subregions: in dorsal V1, vertex pairs with alike ocular polarity exhibited stronger rs-FC than those with unalike ocular polarity, whereas in ventral V1, this trend was reversed—rs-FC was stronger between vertex pairs with unalike ocular polarity than between those with alike ocular polarity.

**Figure 6.**
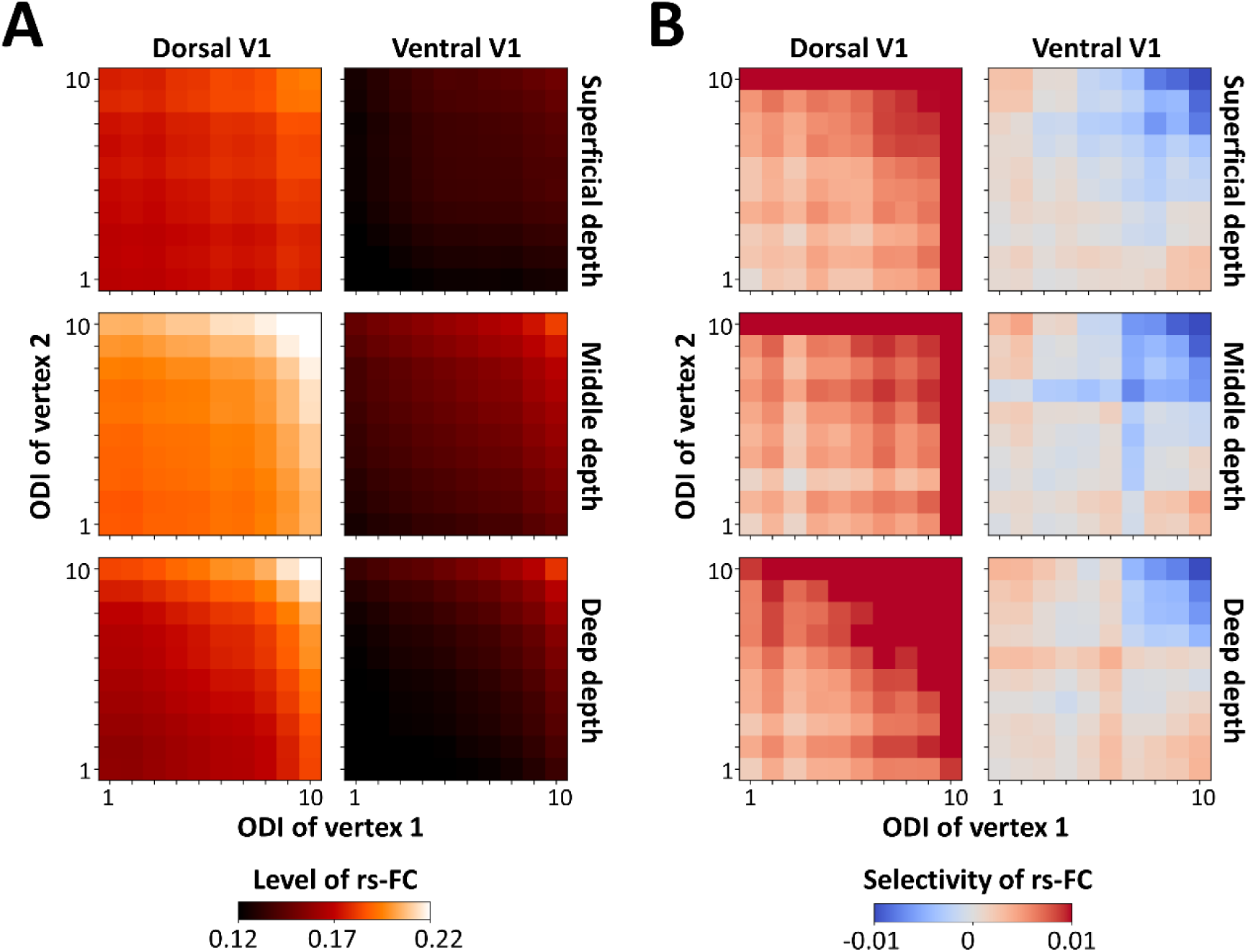
Interhemispheric resting-state functional connectivity (rs-FC) and its selectivity for vertex pairs located within dorsal and ventral primary visual cortex (V1) subregions averaged across all participants. **A)** Similar to the intrahemispheric rs-FC, the interhemispheric rs-FC was higher in the dorsal compared to ventral V1 subregion. The effects of cortical depth and ocular dominance index (ODI) that were observed for intrahemispheric rs-FC are preserved for interhemispheric rs-FC. **B)** In the dorsal V1 subregion, the selectivity of rs-FC was stronger and in favor of pairs with alike ocular polarity. Whereas in the ventral V1 subregions, the selectivity appeared to be weaker and reversed (i.e. in favor of vertex pairs with unalike ocular polarity). The other details are similar to Figure 4.

Consistently, a four-way RM-ANOVA (Supplementary Table 5) revealed a significant main effect of ROI (F(1, 11)=31.89, *p*<10^-3^), a significant interaction between ROI and relative ocular polarity (F(1,11)=6.67, *p*=0.03) and significant three-way interactions between the effects of ROI, relative ocular polarity, and ODI (F(4, 44)=5.35, *p*=0.02) as well as between ROI, ODI and cortical depth (F(8, 88)=3.16, *p*=0.04).

Heterogeneous rs-FC was also observed when comparing interhemispheric rs-FC between central and peripheral V1 subregions. Specifically, again similar to intrahemispheric rs-FC, the relationship between rs-FC and cortical depth varied between central and peripheral regions (Figure 7A). Moreover, the overall pattern of rs-FC selectivity also differed— in the peripheral subregion, selectivity was more pronounced in superficial depths, favoring vertex pairs with alike ocular polarity (Figure 7B). In contrast, in the central subregion, we observed a weaker but opposite trend, with a slight preference for ODCs with unalike ocular polarity. A four-way RM-ANOVA on the measured rs-FC (Supplementary Table 6) revealed a significant interaction between the effect of ROI and cortical depth (F(2, 22)=24.77, *p*<10^-3^), as well as a significant three-way interaction between the effect of ROI, relative ocular polarity, and cortical depth (F(2, 22)=4.74, *p*=0.02) and four-way interaction between all factors (F(8,88)=3.03; *p*=0.048).

**Figure 7.**
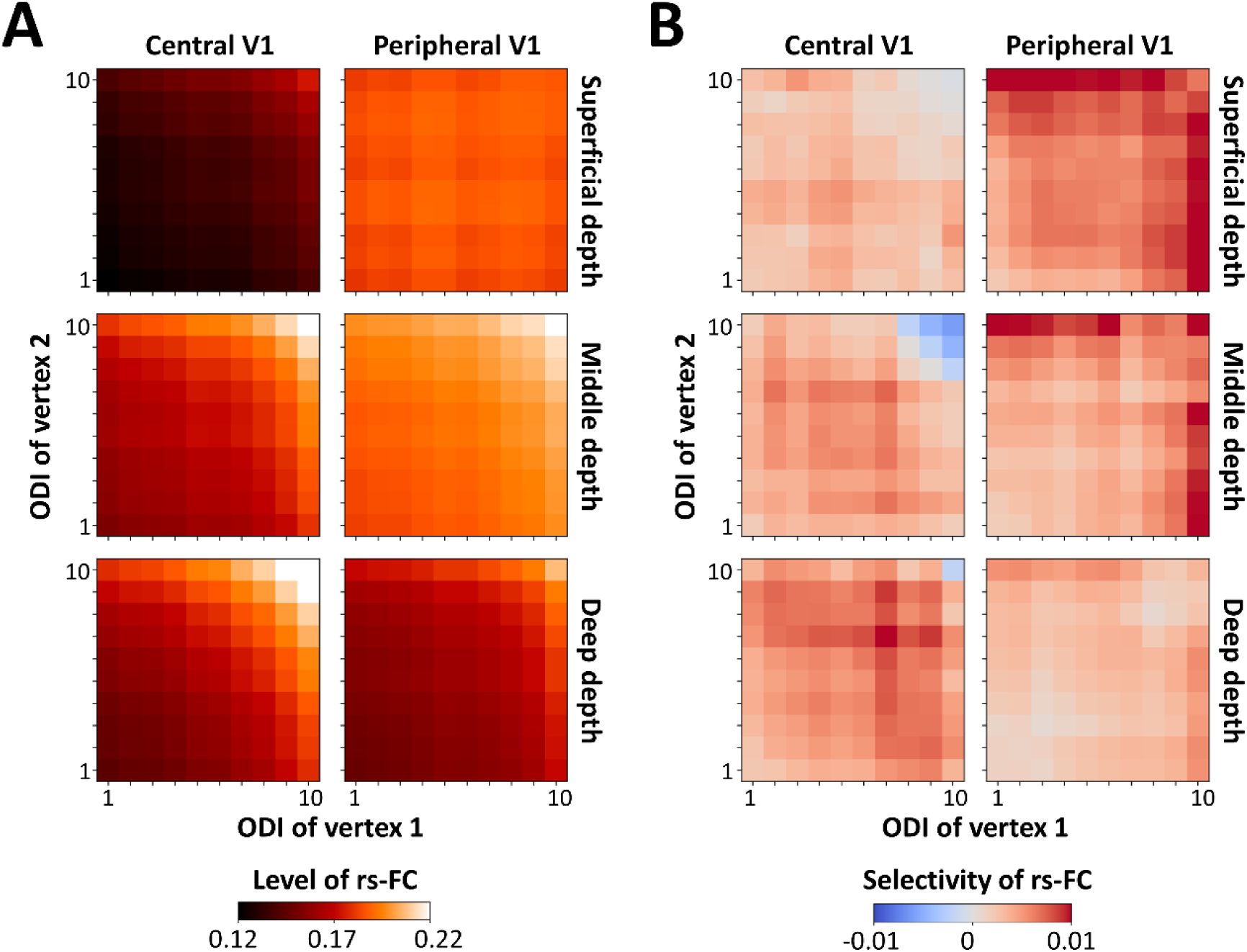
Interhemispheric resting-state functional connectivity (rs-FC) and its selectivity for vertex pairs located within central and peripheral primary visual cortex (V1) subregions averaged across all participants. **A)** The level of interhemispheric rs-FC was, on average, higher in the peripheral compared to the central V1 subregion. Here again, this phenomenon resembles what we detected for the intrahemispheric rs-FC (Figure 5). **B)** The selectivity of rs-FC was higher in peripheral compared to central V1 subregions mainly in superficial and middle cortical depths. The other details are similar to Figure 4.

These findings demonstrate that, like intrahemispheric rs-FC, interhemispheric V1 rs-FC is not uniform but instead varies systematically across V1 subregions, aligning with their specialized roles in visual processing.

### 2.6. Relating ODC maps to rs-FC

Our findings suggest that for each vertex, the relative ocular polarity and the ODI of surrounding vertices may be related to (and even predictable based on) corresponding rs-FC measures. To rigorously test this hypothesis (H1), we pseudo-randomly selected 1,000 seed vertices (10.91% ± 1.76% of all vertices within our V1 ROI) in each hemisphere (see Section 5.5.), generated their 2D rs-FC map with the other vertices, and quantified the correlation between the resultant rs-FC map and the ODC map centered around the seed point (Figure 8A). To mitigate the influence of the fMRI signal’s point spread function (PSF), we excluded the inner 3 mm radius from our analysis. We varied the outer radius from 4 to 10 mm in 0.5 mm increments and compared the measured correlation values against a stringent baseline, defined as the correlation between the same rs-FC pattern and a randomly selected, non-overlapping portion of the ODC map (H0_a_). These analyses were conducted separately for rs-FC measured within deep, middle, and superficial cortical depths.

**Figure 8.**
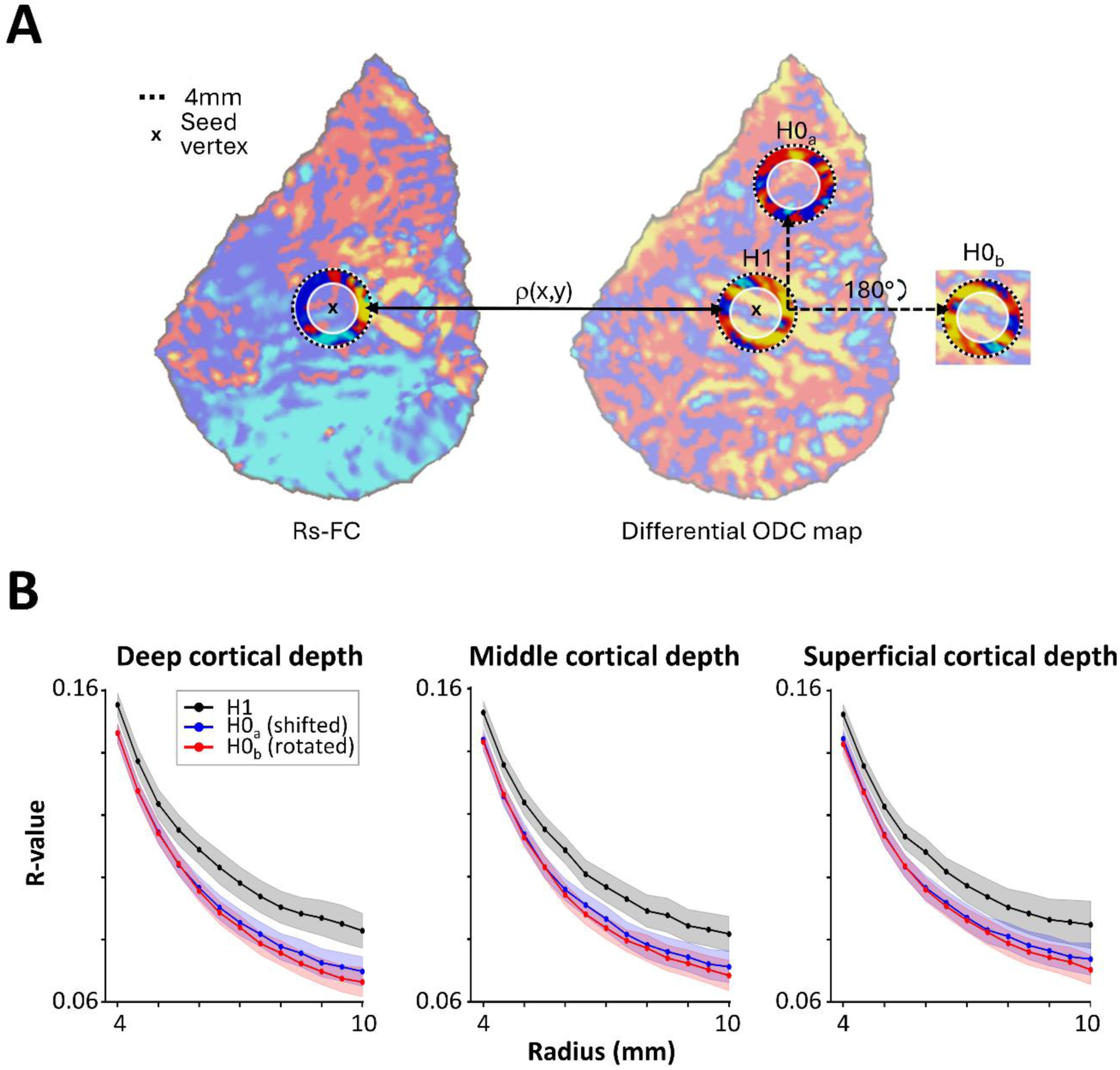
Correlation of resting-state functional connectivity (rs-FC) with the differential ocular dominance column (ODC) map. **A)** Schematic figure showing a map of rs-FC obtained by correlating the time-series of one seed vertex (x) with all other vertices (left) and the corresponding differential ODC map of that representative subject and hemisphere (right). Ring-shaped regions of the rs-FC map with varying outer radii and fixed inner radius of 3mm (white shaded inner circle, to exclude vertex pairs within the fMRI point spread function) were correlated with the ODC differential map at either the ring at the same location (H1) or a shifted location (H0_a_) or the same location but rotated by 180° (H0_b_). **B)** Resulting *r*-values (*y*-axis) averaged across participants are plotted as a function of the ring radius (*x*-axis) for the three different comparisons using rs-FC sampled at the three different cortical depths. The shaded areas correspond to the respective standard error of the mean. For all cortical depths and radii, *r*-values of the H1 were significantly higher than for the two null hypotheses (H0) indicating that the correlation is not only influenced by the overall topography of ODC patterns.

As shown in Figure 8B, although the correlation between ODC and rs-FC maps decreased as the outer radius expanded, it remained higher than either baseline measure across all cortical depths. A three-way RM-ANOVA (measurement type (H1 vs. H0_a_), outer radius, cortical depth), applied to the measured correlation values yielded a significant effect of outer radius (F(12,132)=666.77, *p*<10^-13^) and measurement type (F(1, 11)=15.74, *p*<0.01), with no interaction between the two effects (*p=*0.13). There was no significant main effect of cortical depth (*p*=0.31) or significant interaction between the effect of cortical depth and the other two independent factors (*p*>0.29), indicating that the predictive relationship between rs-FC and ODC organization holds across different cortical depths.

It has already been shown that the organization of ODCs may vary based on eccentricity in the visual field^39^. Given this possibility, it could be argued that the change in the location of the baseline disc could affect the measured correlation values, depending on the distance relative to the actual disc. To rule out this possibility, we conducted a control analysis using an alternative baseline in which the ODC map was only rotated 180° before computing its correlation with the rs-FC pattern (H0_b_). In this condition, the ODC and rs-FC rings remain overlapping relative to each other. Even under this control condition, an RM-ANOVA confirmed robust effects of outer radius (F(12,132)=856.01, *p*<10^-14^), measurement type (F(1, 11)=92.36, *p*<10^-5^), and the significant interaction between the effects of outer radius and measurement type (F(12, 132)=11.60), *p*<10^-4^). The cortical depth showed no significant main or interaction effect with the other factors (*p*>0.12), reinforcing the robustness of our findings across cortical depths.

These results provide empirical support for the hypothesis that rs-FC patterns contain specific information about ODC organization, despite being derived from independent datasets collected in separate scan sessions. The persistence of this relationship across cortical depths and under rigorous control conditions underscores the fundamental link between rs-FC and the underlying columnar architecture of V1.

## 3. Discussion

Rs-FC has been extensively used to investigate functional connections between predefined macroscale brain regions, primarily at the spatial scale of brain areas^40,41^, or to segment the brain into multiple co-fluctuating regions^42–44^. In this study, we provide compelling functional evidence for the selective rs-FC between ODCs — the building blocks of neuronal processing in V1. Our findings reveal key aspects of the mesoscale functional organization of V1, emphasizing heterogeneities across cortical depth and subregions, and offer a fresh perspective on its interhemispheric functional connectivity. Furthermore, we demonstrate new evidence for the predictability of ODC maps based on their rs-FC.

### 3.1. Consistency between rs-FC pattern and anatomical connections

Our findings provide some of the first functional evidence of significantly stronger rs-FC between ODCs with alike compared to those with unalike ocular polarity, in humans, aligning with prior animal studies using postmortem histological methods^4,7,8^ as well as optical imaging and electrophysiological recordings^9–12^. Furthermore, the observation that rs-FC is strongest in middle cortical layers—compared to deep and superficial layers—closely aligns with neuroanatomical studies that demonstrated the highest prevalence of horizontal fibers in layer 4B^6^.

Despite the methodological differences between rs-FC and histological techniques based on using tracers, the overall level of rs-FC, and the level of rs-FC selectivity reported here (Figure 2B) is also comparable to the quantitative measure of this selectivity based on anatomical techniques. Specifically, three previous studies have reported 4%^4^, 8%^8^, and 15%^7^ difference in the number of horizontal fibers connecting ODCs with alike rather than unalike relative ocular polarity. Given that at least two of these studies used the same tracer (biocytin) to detect the horizontal fibers^4,7^, the difference in the level of selectivity can at least be partly attributed to between-species (macaque nemestrina vs. macaque fascicularis) and between-individual variability.

Beyond these functional parallels to structural measurements, our results highlight a strong influence of ODI on rs-FC strength. This effect was robustly detected despite rs-FC and ODI being measured during separate scans collected on different days. Notably, while the previous anatomical studies suggested that the selectivity of horizontal connections is higher in sites that show a stronger ODI^4,7,8^, to our knowledge, the link between ODI and the level of rs-FC has never been explicitly tested in animal studies.

### 3.2. The impact of common input on rs-FC

To minimize the impact of visual input on rs-FC, we instructed the participants to keep their eyes closed during resting-state fMRI scans. This approach significantly reduces the impact of common sensory input, which is known to shape rs-FC patterns^45^, particularly in response to structured visual stimuli^11,46–49^.

Nevertheless, one could still argue that relayed spontaneous activity into V1 via feedforward connections from the lateral geniculate nucleus (LGN) might still contribute to the selective rs-FC observed between ODCs with similar ocular polarity.

Although input from the LGN may play a role in generating spontaneous activity within V1, this input follows a strict visuotopic organization^50^. However, given that rs-FC selectivity was robustly detected over large cortical distances—well beyond the receptive fields of individual V1 columns—the LGN is unlikely to be the driver of the observed phenomenon^9–11^.

Moreover, to mitigate spatial blurring due to the fMRI PSF, we excluded vertex pairs closer than 3 mm, thereby further reducing the potential impact of LGN-driven signal correlations. These methodological choices strengthen the interpretation that the rs-FC structure we observed is not trivially inherited from early feedforward input. Instead, horizontal intracortical connections, are well-positioned to support long-range synchrony within V1^51^.

In parallel, feedback projections from higher-order visual areas could reinforce or modulate these intrinsic patterns^52^, contributing additional structure to the resting-state signal based on learned or experience-dependent priors. However, considering the weak ocular preference of cortical column in extrastriate visual areas^53^, it is unlikely that the feedback from those regions is the main driver of the selective rs-FC between V1 ODCs.

### 3.3. Contribution of polysynaptic connectivity and synchronous neuronal activity to rs-FC

Given these similarities, it is compelling to link the observed rs-FC patterns to the organization of horizontal fibers. While the spatial organization of rs-FC is thought to be at least partially shaped by underlying anatomical connections^41,54–56^, rs-FC is not merely a passive reflection of monosynaptic anatomical connectivity^57^. Instead, the existence of polysynaptic connections and synchronous neuronal activity across ODCs play crucial roles in shaping rs-FC.

The role of polysynaptic connections in generating rs-FC between sites that are not directly connected has been well established^58–61^. This influence is also evident in our findings, which reveal rs-FC between vertex pairs separated by distances of up to 35 mm (Figure 2), far exceeding the reach of a monosynaptic horizontal connection of up to 8mm^1,4,5^. Crucially, our results provide direct evidence that rs-FC selectivity is preserved even across polysynaptic pathways.

Beyond structural connectivity, mounting evidence suggests that rs-FC strength depends on the amplitude of synchronous gamma-band neural activity between the seeded and targeted sites^62–67^. Notably, the first direct evidence of such synchronous activity came from simultaneous electrode recordings from V1 columns that share ocular^9–11^ and orientation^10,11,48,68^ preference—even when these columns are spatially distant and have non-overlapping receptive fields.

Together, these results strongly support the hypothesis that polysynaptic connections within the visual cortex play a crucial role beyond local visual processing. These connections may be instrumental in integrating information across a large visual field, contributing to a more coherent visual perception.

### 3.4. Heterogeneity

Previous studies have shown that global stimulus configuration is more prominent in the lower than in the upper visual field^29,32,69^. Neuroimaging research further reveals that dorsal cortical regions, representing the lower visual field, exhibit heightened sensitivity to visual features critical for global configuration encoding, such as lower spatial frequencies^30^.

Extending these findings, we showed that rs-FC is significantly stronger in dorsal compared to ventral V1 subregions and in peripheral compared to central V1 subregions. Notably, these effects hold for both intrahemispheric and interhemispheric rs-FC (see also Section 3.5.). This pattern aligns with the well-established higher sensitivity for lower spatial frequencies in dorsal versus ventral^31,70,71^ and in peripheral vs. central V1 subregions^33–35^.

Importantly, these findings cannot be explained by variations in gradient-echo BOLD signal intensity across these regions. To clarify, the BOLD signal is generally stronger in the superficial compared to middle and deep cortical depths (Figure 1C) and in central compared to peripheral V1 regions. Yet, rs-FC was strongest in the middle cortical depths and in peripheral V1 subregions, indicating that the observed connectivity patterns are not merely a byproduct of BOLD signal strength. Instead, these results highlight the fundamental role of V1 functional architecture in shaping rs-FC, independent of signal intensity biases.

While our findings on the overall rs-FC pattern across V1 subregions align with the well-established heterogeneity in visual processing across visual fields, their direct link to visual perception remains uncertain without behavioral measurements. One promising direction is to investigate whether individual differences in mesoscale connectivity are predictive of behavioral performance. For instance, reduced connectivity between ODCs might be associated with poorer visual outcomes, such as diminished contrast sensitivity or stereoacuity. Consistent with this hypothesis, studies in animals have shown that strabismus may alter the overall patterns of functional connectivity^8,51^ even though there is still a lack of analogous data in humans (but see^72^ for preliminary findings). Such findings highlight the potential of rs-fMRI as a non-invasive marker of individual visual function and open avenues for future research into clinical populations with visual impairments.

### 3.5. Interhemispheric connections and their role

Interhemispheric rs-FC is crucial for spatial integration around the vertical meridian. Despite its fundamental role in visual perception, our understanding of V1 interhemispheric FC remains limited. This gap in knowledge stems from earlier findings that, compared to downstream areas like V2, the callosal fibers linking the contralateral V1s are sparse and largely confined to a narrow zone along the V1-V2 border^36–38^. Given this and considering that V1 receives extensive feedback from V2—a binocular visual area with robust interhemispheric connections—we assumed that interhemispheric rs-FC in V1 should lack selectivity for ODCs.

Our results overturn this assumption. We demonstrate that, particularly in more peripheral and dorsal regions, rs-FC remains significantly stronger between ODCs with alike rather than unalike ocular polarity. This raises two key possibilities: first, despite their limited numbers, V1 callosal connections may still facilitate selective rs-FC between ODCs in opposite hemispheres. Second, the pronounced rs-FC selectivity in superficial cortical layers is consistent with the hypothesis that indirect pathways, mediated by feedback connections from callosally-connected downstream areas, contribute to this phenomenon.

Yet, it remains unclear how the eye-of-origin information persists within an indirect, polysynaptic network through binocular visual areas. Resolving this will be essential for understanding the mechanisms underlying interhemispheric functional organization in V1.

### 3.6. Mapping ODCs based on rs-FC

Our findings demonstrate that the organization of ODCs and ODI have a direct influence on the pattern of rs-FC across V1. We also provide evidence suggesting a significant relationship between ODC maps and the rs-FC patterns, making a substantial contribution toward leveraging rs-FC for potential segmentation of the visual cortex at mesoscale levels^12^ (see also^20^ for a similar approach outside the visual cortex). To validate the specificity of this correspondence, we compared it to correlations with spatially shifted and rotated ODC maps, which preserve the stripy layout and offer a stricter and more interpretable baseline than a randomized ODC map.

This said, ODCs alone are unlikely to drive the rs-FC pattern. Prior studies in NHPs suggest that orientation and wavelength (color) preference also modulate functional connectivity between cortical columns^10–12,68,73,74^. Consequently, accurate prediction of ODC maps from rs-FC requires disentangling the distinct contributions of these factors on the rs-FC pattern^12^. Future research must refine this approach to fully harness the potential of rs-FC in visual cortex mapping.

### 3.7. Limitations

Although we leveraged an advanced fMRI technique with relatively high spatial resolution, our measurements only indirectly reflect neuronal activity mediated by the BOLD effect and are based on correlations (rather than causal inference). These measurements remain susceptible to partial volume effects and the spatial blurring introduced by the limited spatial resolution and BOLD PSF, which limit the spatial precision of rs-FC estimates^75–77^. Tangentially, we attempted to mitigate PSF-related artifacts by excluding vertex pairs closer than 3 mm in distance, thus reducing the impact of local signal spread. In the radial direction—i.e., across cortical depths—PSF effects are more difficult to constrain, particularly given the overall V1 thickness (∼2mm^78^). This blurring can artificially enhance correlations between layers and obscure depth-specific connectivity patterns. Thus, the true depth-specific differences may in fact be larger than reported, as radial PSF and partial voluming likely attenuate these effects. Future studies employing higher spatial resolution or advanced deconvolution methods may help to better isolate radial connectivity patterns and provide more accurate estimates of depth-specific rs-FC^79,80^.

Additionally, the inherent sensitivity of our fMRI method to pial veins results in stronger yet partially blurred signals in more superficial cortical depths^16,81–83^. In addition to the potential influence of pial veins, intracortical vasculature has been highlighted as a relevant factor in mesoscale fMRI analyses^84^ .

While vascular architecture can indeed introduce spatially structured signal correlations, intracortical vessels generally do not follow the spatial organization of mesoscale neuronal ensembles (e.g. ODCs in V1)^85^. Therefore, while we cannot fully exclude their influence, it is unlikely that they would selectively bias rs-FC between ocular dominance columns with alike eye preference. The observed selectivity in our results is more plausibly attributed to underlying neuronal connectivity rather than shared vascular dynamics.

## 4. Conclusion

The introduction of higher-performance MRI scanners and technological advancements in hardware and software design enable us to now resolve neuronal responses at the fine spatial scale of cortical columns. This study demonstrates the power of high-resolution fMRI in capturing one of the most fine-grained features of cortical organization including the selective connectivity between ODCs. Given that this selectivity produces only a modest 4% to 15% difference in horizontal fiber density, our findings underscore the remarkable sensitivity of high-resolution fMRI in revealing the mesoscale functional organization of the human visual system. Thanks to the short duration of a resting-state scan session (<30 mins) compared to the time necessary for localizing ODCs based on dichoptic visual stimulation (∼3-4 hours^23^) and the absence of cognitive load on the participant, the mesoscale rs-FC measurements present a potentially powerful tool for clinicians to assess visual system integrity at the mesoscale level. It may even enable such studies in patients where conventional methods of ODC mapping are impractical or impossible (e.g., in monocularly blind individuals).

## 5. Methods

### 5.1. Participants

12 subjects (4 females), aged 25-45 years, participated in this study. All participants had radiologically intact brains, intact vision, and no history of neuropsychological disorders. All experimental procedures conformed to the guidelines of the National Institutes of Health (NIH) and were approved by the Mass General Brigham Institutional Review Board. Written informed consent was obtained from all participants prior to all experiments.

### 5.2. Procedure

Each participant was scanned in an ultra-high-field (7T) MRI scanner to collect resting-state data. To test the reproducibility of the findings, four of the participants—selected based on their availability—were rescanned on a different day using a different 7T scanner. For each participant, ODC and retinotopic maps were obtained from a separate set of scans^23^. Structural scans were acquired on a 3T scanner.

#### 5.2.1. Resting-state measurements

To collect spontaneous fMRI activity, each participant was scanned for 3 to 4 runs (512 s per run), except for one individual who was scanned for just one run due to time limitations. Participants were instructed to keep their eyes closed for the duration of the run but not to fall asleep. We communicated with participants between runs to ensure they remained awake during the scan.

#### 5.2.2. ODC mapping

ODC mapping was conducted on different days from the resting-state session. Details of the procedure are already described comprehensively in our recent publication^23^. Briefly, the ODC map was determined by contrasting the fMRI activity evoked in response to stimulation of the dominant eye (DE) vs. non-dominant eye (NDE) in a block design. The stimuli consisted of sparse (5%) horizontally moving random dots (-0.22° to 0.22°; 0.3 Hz), with red (50% of blocks) and green (remaining blocks) dots (0.09° × 0.09°; 56 cd/m²), presented against a black background. Participants viewed the stimuli through custom-made anaglyph spectacles (with red and green filters) mounted on the head coil. The stimuli spanned 20° × 26° of the visual field. Filter laterality (i.e., red-left vs. red-right) was counterbalanced between sessions and across participants.

#### 5.2.3. Retinotopic mapping

The borders of retinotopic visual areas were defined by mapping the horizontal vs. vertical meridians^86,87^. Stimuli consisted of flashing radial checkerboards within retinotopically defined wedge-shaped apertures, extending radially along either the horizontal or vertical meridians (polar angle = 30°) and presented against a gray background. Further details are provided in our previous publication^23^.

### 5.3. Imaging

#### 5.3.1. Main experiments

For the main resting-state fMRI measurements (but see also the reproducibility tests), and also for ODC and retinotopic mapping, we utilized a 7T whole-body Siemens scanner equipped with SC72 body gradients (70 mT/m gradient strength and max slew rate of 200 T/m/s), a custom-built 32-channel helmet radio-frequency (RF) receive coil array, and a birdcage volume RF transmit coil. Resting-state data were collected using single-shot 2D gradient-echo EPI, with 1.0 mm isotropic nominal voxel size, and the following protocol parameter values: TR=4000 ms, TE=28 ms, excitation flip angle=85°, field of view (FOV) matrix=192×192, BW=1184 Hz/pixel, echo-spacing=1 ms, 7/8 phase partial Fourier, 59 oblique-coronal slices covering occipital lobe, acceleration factor R=4 with GRAPPA reconstruction and FLEET- ACS data^88^ with a 10° flip angle. For ODC and retinotopic mapping, we used a similar EPI protocol with TR=3000 ms, flip angle=78°, and 44 oblique-coronal slices.

#### 5.3.2. Reproducibility tests

To test the reproducibility of the resting-state results, 4 participants were scanned again (on a different day) in a Siemens MAGNETOM Terra whole-body MRI (80 mT/m gradient strength and max slew rate of 200 T/m/s). In contrast to the other 7T scanner (see above), this scanner provides passive and active shimming up to 3^rd^ order and is equipped with an in-house-built head RF coil system that use a quadrature birdcage transmit coil along with a close-fitting 64-ch RF receive array^89^. In these sessions, we used single-shot 2D gradient-echo EPI, with 1.0 mm isotropic voxel size, and the following protocol parameter values: TR=4000 ms, TE=25 ms, flip angle=80°, FOV matrix=180×180, BW=1158 Hz/Px, echo-spacing=1 ms, 7/8 phase partial Fourier, 66 oblique-coronal slices covering occipital cortex, and FLEET- ACS data^88^ with 10° flip angle. We also used R=4 dual-polarity GRAPPA reconstruction for improved Nyquist ghost correction^90^.

#### 5.3.3. Structural scans

At ultra-high field (7T), T1-weighted anatomical brain images typically suffer from image contrast variations (bias field) over the FOV due to the combined effects of: (a) transmit RF field inhomogeneity caused by dielectric wavelength effects within the head^91^ and (b) disrupted inversion pulses in areas where tissue susceptibility perturbs the static magnetic field^92^. To avoid this issue, we used anatomical images acquired at 3T on a Siemens TimTrio whole-body scanner, equipped with the standard vendor-supplied 32-channel RF head coil array, which provides relatively uniform image contrast over the full FOV. We used a 3D T1-weighted MPRAGE sequence with protocol parameter values: TR=2530 ms, TE=3.39 ms, TI=1100 ms, flip angle=7°, bandwidth=200 Hz/pix, echo spacing=8.2 ms, voxel size=1.0×1.0×1.0 mm³, and FOV=256×256×170 mm³.

### 5.4. Data analysis

Functional and anatomical MRI data were pre-processed and analyzed using FreeSurfer (version 7.4.0; http://surfer.nmr.mgh.harvard.edu)^93^ and in-house MATLAB and Python code. As described in the following sections in detail, each subject was analyzed entirely in their native anatomical space to preserve the fine-grained structure of ODCs and cortical lamination. This native-space processing was maintained throughout all individual analyses. An overview of the processing steps applied to the ODC mapping and resting-state data is presented in Supplementary Figure 4.

#### 5.4.1. Anatomical data

##### 5.4.1.1. Defining cortical surfaces

Based on each participant’s anatomical data, cortical surfaces were reconstructed using FreeSurfer’s recon-all pipeline, which includes intensity normalization, skull stripping, white and pial surface extraction, and topological correction using a deformable surface model constrained by neuroanatomical priors^94^. In this process, the standard pial surface was defined as the boundary between the gray matter (GM) and the surrounding cerebrospinal fluid (CSF), while the white matter (WM) surface was generated as the interface between the WM and GM. Additionally, nine intermediate surfaces were created using the equivolume approach, spaced at intervals of 10% of the cortical thickness. Finally, surfaces were flattened using FreeSurfer’s mris_flatten routine.

##### 5.4.1.2. Defining cortical depths

To examine the impact of cortical depth on rs-FC, the GM was divided into deep, middle, and superficial cortical depths (Figure 1A). The deep cortical depth was defined as the region between the WM-GM interface and the two adjacent surfaces above it. The superficial cortical depth was defined as the region between the GM-CSF interface and the two adjacent surfaces below it. The middle cortical depth included the three intermediate reconstructed surfaces. Notably, this approach left one intermediate surface gap between these depth levels to improve the separation of the three segments. To enhance the co-registration of functional and structural scans, all surfaces were upsampled to a 0.5 mm resolution using butterfly subdivision in mris_mesh_subdivide as implemented in FreeSurfer and outlined in our previously introduced pipeline^95^.

#### 5.4.2. Functional data

##### 5.4.2.1. Preprocessing

All functional data were preprocessed with a pipeline for ultra-high-resolution fMRI which mostly uses FreeSurfer commands^95^. Briefly, the collected functional data were first upsampled to 0.5 mm isotropic resolution using trilinear interpolation using FreeSurfer’s mri_convert (see also^95^). For each participant, functional data from each run were rigidly aligned (with 6 degrees of freedom) to their structural scan using rigid Boundary-Based Registration (Supplementary Figure 5)^96^. Head motion covariates were then derived for six directions. After motion correction, intensity normalization was applied to the fMRI data collected during ODC mapping, but not to the resting-state fMRI data (Supplementary Figure 4).

Resting-state data were analyzed with an adjusted preprocessing pipeline designed to preserve as much of the original signal as possible while minimizing the impact of noise sources, which is particularly important when examining mesoscale functional connectivity. Specifically, before upsampling to 0.5 mm, the data were denoised using the NORDIC approach^97^, which was reported to effectively suppresses thermal noise in fMRI data^98^. Thermal noise poses a particular challenge for resting-state analyses, as it reduces temporal signal-to-noise ratio (tSNR) and can mask meaningful low-frequency BOLD fluctuations, thereby compromising the reliability of functional connectivity estimates. MP-PCA denoising, as implemented in NORDIC, has been shown to approximately double the tSNR and substantially increase correlation coefficients, with only a minimal increase in spatial smoothness (7%)^99^. After slice timing correction^93^, the data were detrended for quadratic trends and high-pass filtered using a cut-off frequency of 0.01 Hz ^100^.

To preserve spatial resolution on the surface, no tangential spatial smoothing was applied to the fMRI data. Instead, radial (intracortical) smoothing—perpendicular to the cortex and within cortical columns— was used^101^. Considering the cortical thickness of V1 (∼2mm^78^), the 1 mm isotropic voxel size, and the fact that voxel boundaries do not precisely align with the gray–white matter interface, each cortical depth level likely receives signal contributions from approximately 1–2 voxels. For ODC and retinotopic mapping, to minimize the blurring impact of pial veins on functional map resolution^16,81–83^, radial smoothing was restricted to the deep cortical depth (as defined in 5.4.1.2). For rs-FC analysis, radial smoothing was applied either to the entire GM (sections 2.1-2.2) or separately to the deep, middle, and superficial cortical depths (sections 2.3-2.6). Notably, previous studies in NHPs ^102^ and humans ^23^ have shown that the ODC maps remain mostly unchanged across cortical depth. The stability and consistency of ODC maps derived from fMRI (across sessions and depth levels) are already demonstrated in our previous publications^16,23^.

Notably, we avoided applying any method of spatial distortion correction to the fMRI data because EPI distortions are typically small in the occipital lobe^103^. This point is also demonstrated in Supplementary Figure 5, which highlights the alignment of EPI to anatomical scans in the occipital lobe, including V1. Moreover, distortion correction techniques generally cause spatial smoothing^95^.

Considering that spatial resolution is critical for resolving fine-scale structures such as ODCs, we avoided such smoothing-inducing corrections.

##### 5.4.2.2. Resting-state functional connectivity analysis

The rs-FC between pairs of vertices within the flattened cortical surface mesh of V1 was measured by estimating Pearson’s partial correlation coefficients of their resting-state fMRI time series. For our main analysis, this was conducted separately within each hemisphere; however, we also evaluated rs-FC between hemispheres. We did not include correlations between vertex pairs located at different cortical depths considering the organization of horizontal fibers (i.e., primarily parallel to the cortical surface).

To reduce the impact of physiological and motion-related confounds, we included the mean white matter signal and six head motion parameters as covariates in the partial correlation analyses. Moreover, to remove the impact of fMRI signal propagation, we excluded all vertex pairs with an inter-vertex distance less than 3 mm. This threshold is in line with prior estimates of the fMRI PSF at 7T^75–77^ and is intended to reduce the influence of non-neuronal spatial autocorrelation. This approach allowed us to estimate functional connectivity while accounting for structured sources of physiological noise. Notably, we avoided using other more recently introduced methods that rely on specific assumptions about the correlated spontaneous activity evoked between voxels because they have not been tested at mesoscale _levels104,105._

To mitigate the influence of noise at the single-vertex level, we averaged correlation coefficients across runs, improving the signal-to-noise ratio. For our next analyses, which required linear statistical operations on correlation values, we used Fisher *z*-transformed correlation values that are expected to have a distribution closer to normal than Pearson’s correlation coefficients do and are thus more suitable for parametric calculations.

##### 5.4.2.3. ODC and retinotopic mapping

Details of the data analysis for mapping ODCs and retinotopic visual areas are described in our recent publication^23^. Briefly, after preprocessing (see above), a standard hemodynamic response model based on a gamma function was fitted to the signals from deep cortical depths to estimate the amplitude of the blood-oxygen-level-dependent (BOLD) response. To generate differential maps of DE vs. NDE activation, a vertex-wise general linear model was applied by computing the contrast of interest. The resulting beta and significance maps were projected onto each participant’s anatomical volumes and reconstructed cortical surfaces (Figure 1B). The reproducibility of the maps across scan sessions was tested and proved for all individual participants^16,23^. As noted above, only data from deep cortical depths were used for ODC and retinotopic mapping.

### 5.5. Statistical analysis

As a second-order statistical analysis (see also Section 5.4.2.2), we assessed the significance of the impact of the following independent parameters on rs-FC: (1) relative ocular polarity, (2) distance between vertices, (3) level of ocular preference (ocular dominance index (ODI)), (4) cortical depth, and (5) region of interest (ROI). These parameters were calculated as follows.

#### 5.5.1. Distance between vertex pairs

The distance between vertices in each pair was calculated as the Euclidean distance on the flattened cortical surface. To minimize the bias in the rs-FC resulting from the effect of the fMRI PSF, we excluded vertex pairs located closely together (<3 mm)^75–77^. Since vertex pairs with alike ocular polarity (see below) generally were closer together than unalike pairs, we used random subsampling to balance the distance distributions between alike and unalike vertex pairs. To that end, we divided the distance distribution of all vertex pairs into 100 quantiles, for each of which, we then determined the counts of alike and unalike vertex pairs and randomly discarded excess pairs from the larger group to equalize their numbers. Because V1 varied in size across participants, vertex pairs for each participant were divided into 10 distance quantiles, with 1 referring to the shortest distances and 10 referring to the longest distance.

The quantile rank (rather than the absolute distance) was used for further statistical analysis. The median distance for each quantile is shown in Supplementary Figure 1. The distance between vertex pairs was then used as a numerical parameter in the statistical tests.

#### 5.5.2. Relative ocular polarity of vertices

To assess the effect of relative ocular polarity on rs-FC, a pair of vertices was classified as alike or unalike when the ocular preferences of the two vertices were the same (i.e. both vertices responded preferentially to the same eye) or opposite, respectively. The relative ocular polarity was then used as a categorical parameter in the statistical tests. Notably, the ocular preference of each vertex was determined based on the unthresholded ODC map (Figure 1B). Consistent with our previous findings in a smaller group of participants^23^, the number of vertices responding preferentially to DE and NDE stimulation, as well as the amplitude of ocular-preferring activity (i.e., the response to the preferring vs. non-preferring eye stimulation), did not differ significantly (Supplementary Figure 2). This result indicates that both eyes were represented equally within V1.

#### 5.5.3. Ocular dominance index (ODI)

To assess the impact of ocular preference level on rs-FC, we used a method similar to the one introduced by Hubel and Wiesel, 1962 and later adopted by others^106–108^. Specifically, for each vertex, the level of ocular preference — i.e., the estimated beta value for differential response to preferred vs. non-preferred eye stimulation —was measured during ODC mapping. For each subject, these beta values were divided into 10 quantiles, with 1 representing the weakest ocular preference and 10 representing the strongest.

The quantile rank (rather than the absolute ocular preference) was then used as the ODI. The median beta value for each ODI is shown in Supplementary Figure 6.

The effect of ODI on rs-FC between vertex pairs was tested by grouping vertices into 5 cumulative categories: (1) vertex pairs from quantiles 1 and 2, (2) quantiles 1 to 4, (3) quantiles 1 to 6, (4) quantiles 1 to 8, and (5) quantiles 1 to 10.

#### 5.5.4. Cortical depths

To test the impact of cortical depth on the level of rs-FC, the rs-FC data analysis was conducted separately for the deep, middle, and superficial depths of the GM. The cortical depth was then used as a categorical parameter in the statistical tests.

#### 5.5.5. ROI

In the main analysis (sections 2.1 to 2.3), the ROI included the stimulated portion of V1 (radius<10°) except for a small dorsal region along the V1-V2 border (Figure 1B) where ODCs could not be identified due to occlusion of the ventral visual field by the nose^23^. To evaluate the heterogeneity of rs-FC within V1, we manually divided the stimulated part of V1 into size-matched dorsal and ventral as well as central (radius<4°) and peripheral (4°<radius<10°) subregions and compared the rs-FC within these subdivisions (see sections 2.4 and 2.5). The ROI was then used as a categorical parameter in the statistical tests.

#### 5.5.6. Statistical tests

##### 5.5.6.1. Repeated measures (RM) ANOVA

RM-ANOVA was primarily used to test the impact of independent parameters (see above) on the measured rs-FC between the vertex pairs. Specifically, we first tested the impact of relative ocular polarity (alike vs. unalike) and distance using a two-way RM-ANOVA (Section 2.1). Second, for a subset of 4 participants, the reproducibility of the results was tested by adding the session number (categorical: 1^st^ vs. 2^nd^) to the independent parameters (i.e., distance and relative ocular polarity) and applying a three-way RM-ANOVA (Section 2.2).

The impact of ODI (5 cumulative categories, see Section 5.5.3) and cortical depth, along with relative ocular polarity, was tested by applying a three-way RM-ANOVA (Section 2.3). Lastly, heterogeneity of intrahemispheric and interhemispheric rs-FC measurements was tested by including V1 subregions (ROIs) in the list of independent parameters and applying a four-way RM-ANOVA (sections 2.3 and 2.4).

Notably, in all tests except for interhemispheric rs-FC measurements, the rs-FC values from the two hemispheres were averaged to improve the signal-to-noise ratio. Considering that RM-ANOVA is particularly susceptible to violations of the sphericity assumption, caused by the correlation between measured values and unequal variance of differences between experimental conditions, results were corrected, when necessary (as determined using a Mauchly test), using the Greenhouse-Geisser method. A p-value of <0.05 was considered statistically significant.

##### 5.5.6.2. Linear mixed effects (LME) model

To ensure that overlap between the cumulative ODI categories did not skew the estimated significance of our results, we conducted a supplementary analysis based on using an LME model to assess the relationship between rs-FC and relative ocular polarity, ODI, and cortical depth accounting for subject-level variability (random intercepts by subject). In this analysis, rather than using the cumulative ODI categories, we used the product of BQ_1_ and BQ_2_ (i.e. BQ_12_), where BQ_1_ and BQ_2_ represent the ODIs of the first and second vertices in a pair. Since this created a symmetric matrix of 10×10 categories, we excluded the redundant upper triangle, resulting in 55 joint ODI categories, which were then used in an LME model (see Section 5.5.6.2). By using 55 non-overlapping joint ODI brackets, we ensured that the significance of the results was not overestimated due to datapoint (vertex pair) interdependencies.

To apply the model the MATLAB command `fitlme` was used according to the following formula: rsFC ∼ BQ_12_ + CD + ROP + BQ_12_ * CD + BQ_12_ * ROP + CD * ROP + BQ_12_ * CD * ROP + (1|Subject), where BQ_12_ represents the product of the ODIs of the first and second vertices in a pair, CD represents cortical depth, and ROP is relative ocular polarity. This approach enabled us to check for the consistency of the results obtained by ANOVA and LME. Notably, in contrast to ANOVA, LME takes the hierarchical structure of the data into account. This method further allowed us to quantify the independent contributions of the main factors and their interactions while controlling for variability across subjects (see Section 2.3). A p-value of <0.05 was considered statistically significant.

#### 5.5.6. Correlation between rs-FC and ODC maps

To evaluate how well rs-FC reflects the underlying ODC map, we conducted a correlation analysis. We selected 1000 seed vertices at random within V1 (far enough from the V1 border) and correlated each seed vertex’s resting-state fMRI time-series with the rest of the vertices in a 2D donut-shaped region (ring) centered at that seed vertex, generating 1000 2D rs-FC maps. To investigate the impact of distance, we conducted the analysis multiple times within concentric rings around each seed vertex with varying outer radius ranging from 4 to 10 mm. Similarly to our main analyses, we excluded nearby target vertices falling within an inner radius of 3 mm to minimize potential biases caused by the PSF of the BOLD signal^75–77^. Within each ring, we calculated the correlation between the rs-FC map and the underlying ODC map, which constitutes the correlation value under the alternative hypothesis (H1) that rs-FC and ODC maps are related. For each ring size, the resultant correlation values were averaged across the 1000 sampled sites.

To determine the statistical significance of our results, we compared these correlations to null-hypothesis (H0) distributions (correlations between unrelated rs-FC and ODC maps) generated by correlating the rs-FC values in each ring with the ODC differential map in a spatially shifted (misaligned) ring (H0_a_) as well as a 180° rotated ring (H0_b_). With this approach, we tested whether the observed correlations were meaningful and not solely due to the inherent spatial structure of the ODC map. To ensure that interpolation and resampling artifacts affected both the null and alternative tests similarly, in all experiments, we resampled the rs-FC and ODC map values with nearest-neighbor interpolation on a set of points generated via 2D quasi-random Halton sampling^109,110^ and distributed with a density of 10 points per mm².

### 5.6. Data availability

The data and code will be shared upon reasonable request.

## Acknowledgment

This work was supported by the National Eye Institute (R01EY030434 and R01EY029713), the National Institute on Aging (RF1AG068261), the Deutsche Forschungsgemeinschaft (DFG, German Research Foundation) – project no. 347592254 (WE 5046/4-2 and/or KI 1337/2-2), the Federal Ministry of Education and Research (BMBF) under support code 01ED2210, as well as by the MGH/HST Athinoula A. Martinos Center for Biomedical Imaging and the Max Planck School of Cognition. Crucial imaging resources were made available by the NIH Shared Instrumentation Grant S10RR019371. We thank Azma Mareyam for helping with hardware maintenance as well as John Kirsch and Daniel Park for their support in raw data export during this study. We thank Amanda Nabasaliza, Sarala N. Malladi, and Bryan Kennedy for their help with recruitment and data collection. We also thank Drs. Antony Morland and Jingyuan Chen for their comments.

## Abbreviations

ODCs: Ocular Dominance Columns
FMRI: Functional Magnetic Resonance Imaging
RS-FC: Resting-state Functional Connectivity
DE: Dominant Eye
NDE: Non-Dominant Eye
ODI: Ocular Dominance Index
ROI: Region Of Interest
RM-ANOVA: Repeated Measures Analysis of Variance

## Author’s contribution

**Conceptualization, formal analysis, visualization, writing – original draft:** M.E.S., I.A. and S.N.

**Investigation, writing – review & editing:** M.E.S., I.A., J.S., B.B., Y.C., W.S.H., E.K., N.W. and S.N.

**Methodology, software:** M.E.S., I.A., J.S., B.B., Y.C., W.S.H. and S.N.

**Data curation, validation:** M.E.S. and S.N. Supervision: E.K., N.W. and S.N.

**Funding acquisition:** N.W. and S.N.

**Resources, project administration:** S.N.

