## Supplementary materials for "Unraveling the mesoscale resting-state functional connectivity of ocular dominance columns in humans using high-resolution functional MRI"

### Supplementary Figures

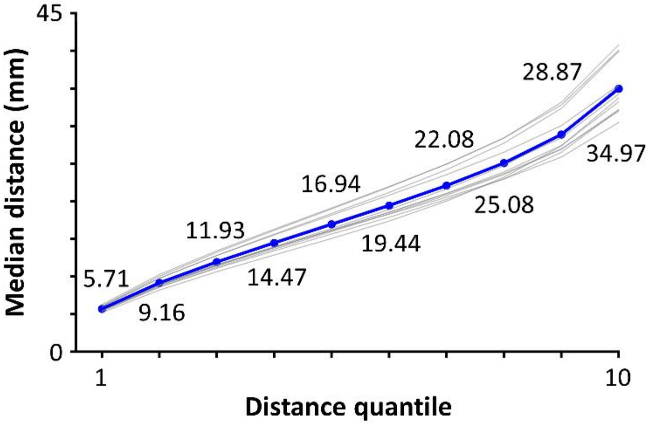


**Supplementary Figure 1.** Median distances for 10 distance quantiles. The mean value and the values from individual subjects are shown by blue bold and gray curves, respectively. The distance quantiles (rather than the absolute distances) were used to account for interindividual differences in the size of V1.

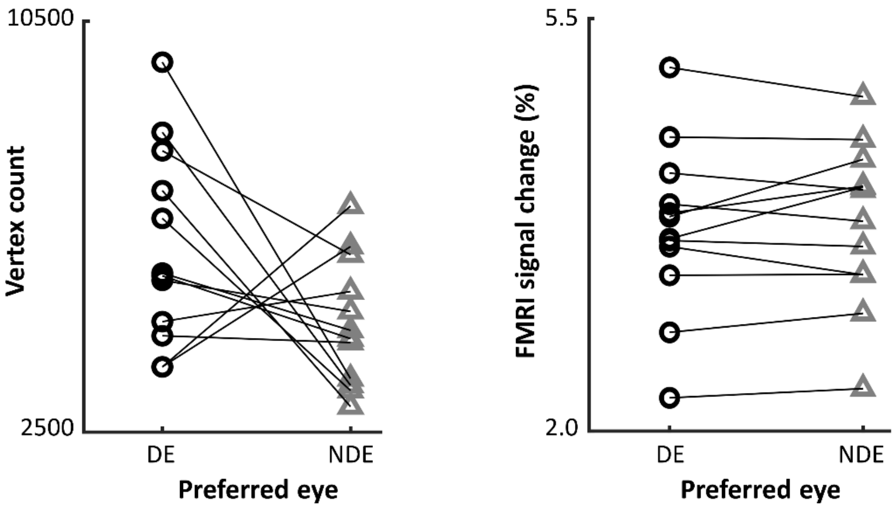


**Supplementary Figure 2.** Number of dominant eye (DE) versus non-dominant eye (NDE) preferring vertices in V1. Vertex count (left) and the level of ocular preference (fMRI percent signal change; right) did not differ significantly between vertices responding to either DE (black circles) or NDE (gray triangles) across participants. Data points of each subject are connected by a line segment.

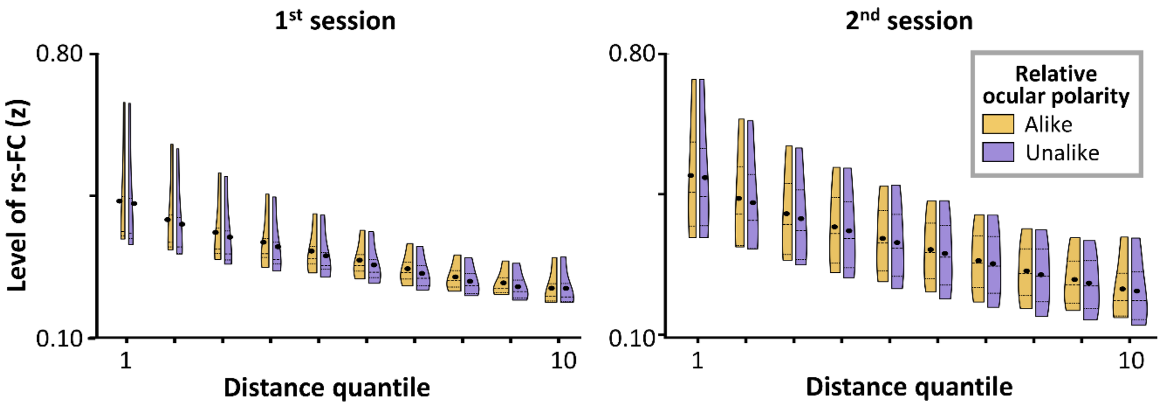


**Supplementary Figure 3.** Reproducibility of resting-state functional connectivity (rs-FC) pattern across different days for 4 individuals. Mean rs-FC levels (*z*-values) across subjects are displayed using split violin plots for vertex pairs with alike (orange) and unalike (violet) ocular polarity for 10 distance quantiles. The effects of relative ocular polarity (i.e., alike>unalike) were apparent in both sessions, and the interaction between the effects of session and relative ocular polarity remained non-significant. The rest of the details are similar to Figure 2A.

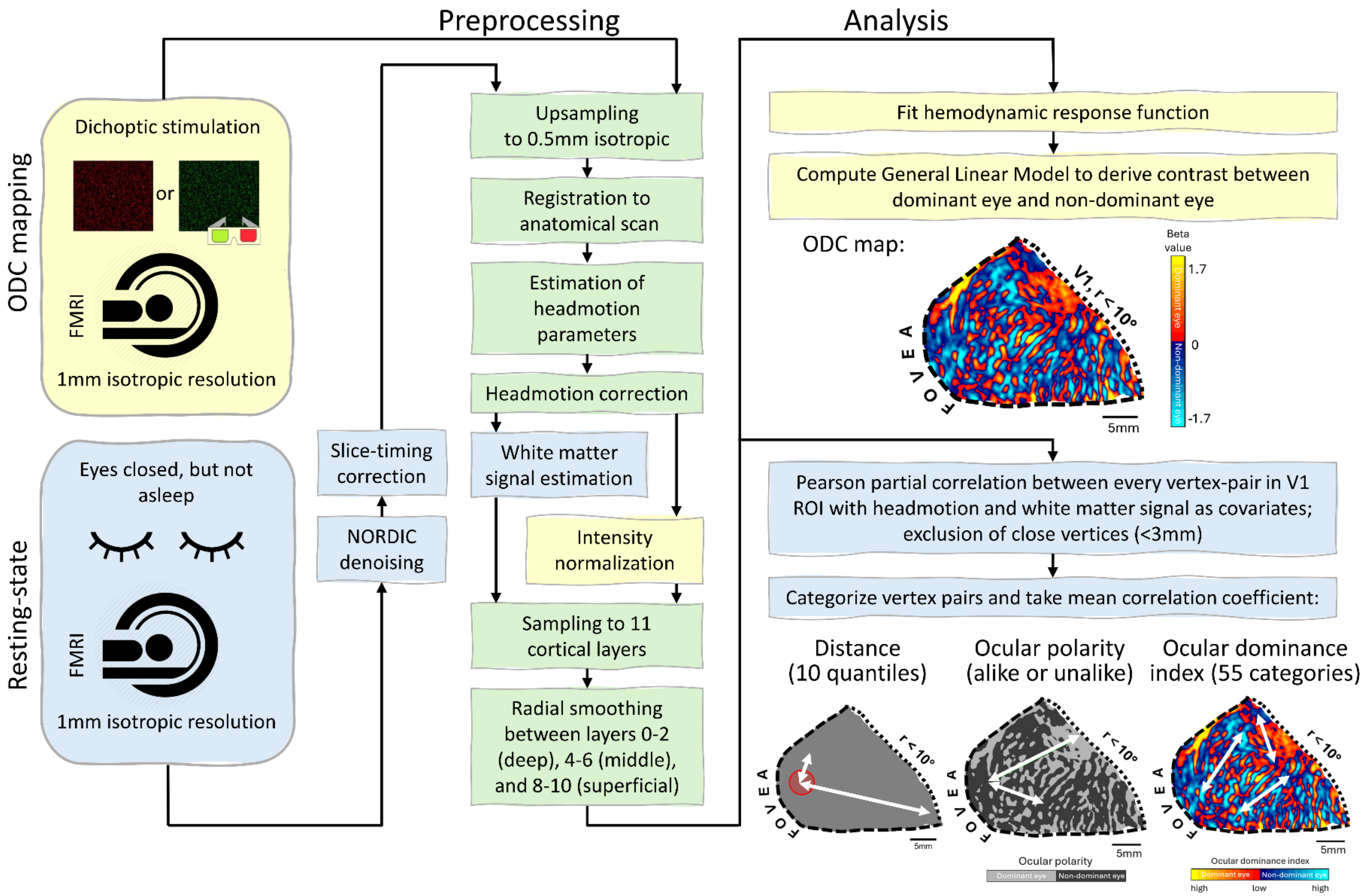


**Supplementary Figure 4**. Graphical overview of the data acquisition, preprocessing, and analysis steps for ocular dominance column mapping (yellow shading) and resting-state fMRI data (blue shading), collected on different days. Shared steps across both modalities are highlighted in green.

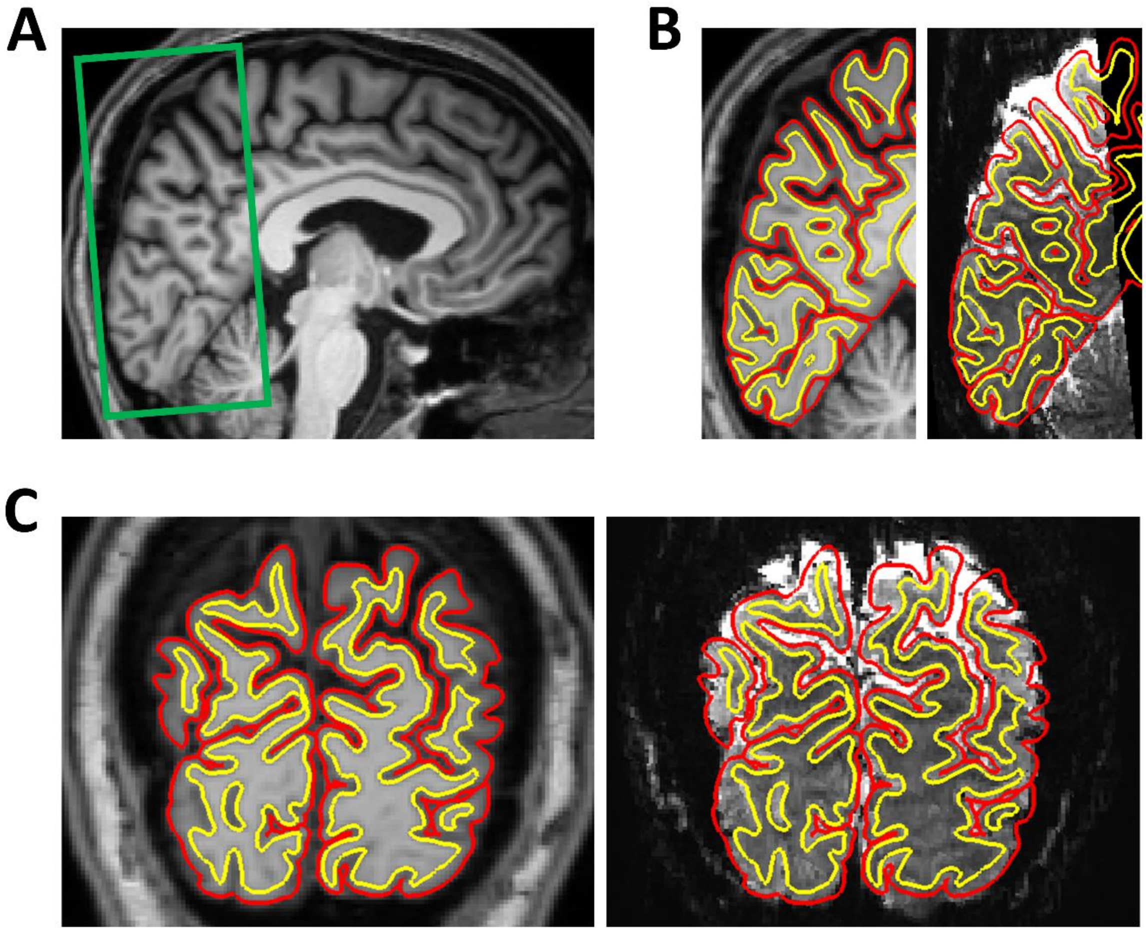


**Supplementary Figure 5.** Co-registration results for one participant. A) Functional field-of-view (FOV) placement (green box) overlaid on anatomical scan. The FOV was placed to maximize coverage of the stimulated portion of V1 (radius<10°). B-C) Alignment of the functional images to their corresponding anatomical scans in sagittal and coronal views, respectively. Red and yellow contour lines indicate the pial surface and the gray–white matter boundary, respectively, defined based on each participant's anatomical reconstruction (left subpanel). The close alignment between the functional signal boundaries and the anatomical surfaces demonstrates accurate co-registration (right subpanel).

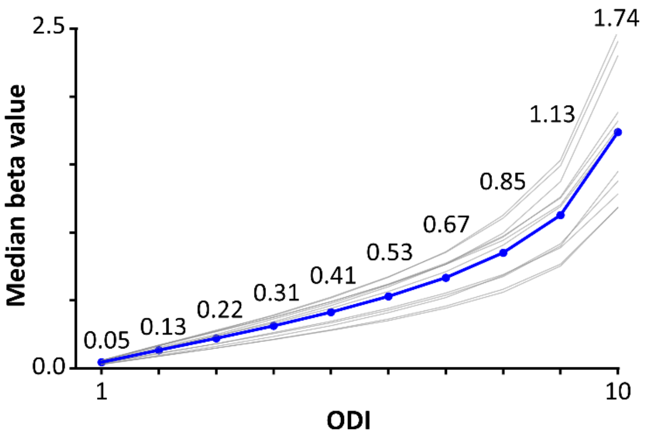


**Supplementary Figure 6.** The level of ocular preference (response to stimulation of preferred – non-preferred eye) for the 10 ocular dominance index (ODI) quantiles. The mean value and the values from individual subjects are shown by blue bold and gray curves, respectively. The variance across subjects in ODI highlights the importance of accounting for interindividual differences by choosing quantiles instead of fixed thresholds. The other details are similar to Supplementary Figure 1.

### Supplementary Tables

**Supplementary Table 1.** Results of applying RM-ANOVA to estimating the effects of ocular dominance index (ODI), relative ocular polarity, and cortical depth on resting-state functional connectivity (rs-FC) within V1.

| ***Effect*** | ***df*** | ***F*** | ***p*** |
|---|---|---|---|
| ***ODI*** | 4, 44 | 11.39 | <0.01** |
| ***Relative ocular polarity*** | 1, 11 | 6.09 | 0.03* |
| ***Cortical depth*** | 2, 22 | 4.74 | 0.047* |
| ***ODI ×<br>relative ocular polarity*** | 4, 44 | 6.08 | 0.02* |
| ***ODI ×<br>cortical depth*** | 8, 88 | 13.47 | $<10^{-4}$*** |
| ***Relative ocular polarity ×<br>cortical depth*** | 2, 22 | 1.25 | 0.30 |
| ***ODI ×<br>relative ocular polarity ×<br>cortical depth*** | 8, 88 | 1.80 | 0.20 |

df = degrees of freedom; *: $p$<0.05; **: $p$<0.01; ***: $p$<0.001

**Supplementary Table 2**. Results of applying a linear mixed effects (LME) model to estimate the effects of ocular dominance index (ODI), relative ocular polarity (unalike relative to alike), and cortical depth (middle and superficial cortical depth relative to deep cortical depth) on resting-state functional connectivity (rs-FC) within V1.

| ***Effect*** | ***t*** | ***p*** |
|---|---|---|
| ***ODI*** | 16.83 | $<10^{-61}$*** |
| ***Unalike ocular polarity*** | 2.00 | 0.045* |
| ***Middle cortical depth*** | 15.14 | $<10^{-50}$*** |
| ***Superficial cortical depth*** | 6.56 | $<10^{-10}$*** |
| ***ODI × unalike ocular polarity*** | 2.67 | <0.01* |
| ***ODI × middle cortical depth*** | -3.62 | $<10^{-3}$*** |
| ***ODI × superficial cortical depth*** | -7.17 | $10^{-12}$*** |
| ***Unalike ocular polarity × middle cortical depth*** | -0.31 | 0.76 |
| ***Unalike ocular polarity × superficial cortical depth*** | -0.39 | 0.69 |
| ***ODI × unalike ocular polarity × middle cortical depth*** | -0.43 | 0.66 |
| ***ODI × unalike ocular polarity × superficial cortical depth*** | -0.73 | 0.47 |

*: $p<0.05$; **: $p<0.01$; ***: $p<0.001$

**Supplementary Table 3**. Results of applying RM-ANOVA to compare the effects of region-of-interest (ROI), relative ocular polarity, ocular dominance index (ODI), and cortical depth on resting-state functional connectivity (rs-FC) in dorsal vs. ventral portions of V1.

| *Effect* | *df* | *F* | *p* |
|---|---|---|---|
| ***ROI*** | 1, 11 | 29.26 | $<10^{-3}$*** |
| ***Relative ocular polarity*** | 1, 11 | 4.13 | 0.07 |
| ***ODI*** | 4, 44 | 11.38 | $<10^{-3}$*** |
| ***Cortical depth*** | 2, 22 | 3.38 | 0.06 |
| ***ROI ×<br>relative ocular polarity*** | 1, 11 | 1.52 | 0.24 |
| ***ROI ×<br>ODI*** | 4, 44 | 2.29 | 0.07 |
| ***ROI ×<br>cortical depth*** | 2, 22 | 0.60 | 0.56 |
| ***Relative ocular polarity ×<br>ODI*** | 4, 44 | 2.05 | 0.17 |
| ***Relative ocular polarity ×<br>cortical depth*** | 2, 22 | 2.13 | 0.17 |
| ***ODI ×<br>cortical depth*** | 8, 88 | 12.69 | $<10^{-4}$*** |
| ***ROI ×<br>relative ocular polarity ×<br>ODI*** | 4, 44 | 1.23 | 0.31 |
| ***ROI ×<br>relative ocular polarity ×<br>cortical depth*** | 2, 22 | 5.17 | 0.03* |
| ***ROI ×<br>ODI ×<br>cortical depth*** | 8, 88 | 4.38 | 0.02* |
| ***Relative ocular polarity ×<br>ODI ×<br>cortical depth*** | 8, 88 | 0.65 | 0.47 |
| ***ROI ×<br>relative ocular polarity ×<br>ODI ×<br>cortical depth*** | 8, 88 | 0.65 | 0.51 |

df = degrees of freedom; *: $p<0.05$; **: $p<0.01$; ***: $p<10^{-3}$

**Supplementary Table 4**. Results of applying RM-ANOVA to compare the effects of region-of-interest (ROI), relative ocular polarity, ocular dominance index (ODI), and cortical depth on resting-state functional connectivity (rs-FC) within central vs. peripheral V1.

| *Effect* | *df* | *F* | *p* |
|---|---|---|---|
| ***ROI*** | 1, 11 | 1.58 | 0.23 |
| ***Relative ocular polarity*** | 1, 11 | 5.52 | 0.04* |
| ***ODI*** | 4, 44 | 16.41 | $<10^{-3}$*** |
| ***Cortical depth*** | 2, 22 | 4.08 | 0.06 |
| ***ROI ×<br>relative ocular polarity*** | 1, 11 | 0.11 | 0.74 |
| ***ROI ×<br>ODI*** | 4, 44 | 3.59 | 0.07 |
| ***ROI ×<br>cortical depth*** | 2, 22 | 27.72 | $<10^{-4}$*** |
| ***Relative ocular polarity ×<br>ODI*** | 4, 44 | 8.15 | <0.01** |
| ***Relative ocular polarity ×<br>cortical depth*** | 2, 22 | 3.27 | 0.08 |
| ***ODI ×<br>cortical depth*** | 8, 88 | 23.81 | $<10^{-6}$*** |
| ***ROI ×<br>relative ocular polarity ×<br>ODI*** | 4, 44 | 0.23 | 0.71 |
| ***ROI ×<br>relative ocular polarity ×<br>cortical depth*** | 2, 22 | 0.06 | 0.88 |
| ***ROI ×<br>ODI ×<br>cortical depth*** | 8, 88 | 0.76 | 0.48 |
| ***Relative ocular polarity ×<br>ODI ×<br>cortical depth*** | 8, 88 | 3.45 | 0.06 |
| ***ROI ×<br>relative ocular polarity ×<br>ODI ×<br>cortical depth*** | 8, 88 | 0.63 | 0.53 |

df = degrees of freedom; *: $p<0.05$; **: $p<0.01$; ***: $p<10^{-3}$

**Supplementary Table 5**. Results of applying RM-ANOVA to compare the effects of region-of-interest (ROI), relative ocular polarity, ocular dominance index (ODI), and cortical depth on interhemispheric resting-state functional connectivity (rs-FC) between dorsal vs. ventral V1 subregions.

| *Effect* | *df* | *F* | *p* |
|---|---|---|---|
| ***ROI*** | 1, 11 | 31.89 | $<10^{-3}$*** |
| ***Relative ocular polarity*** | 1, 11 | 2.73 | 0.13 |
| ***ODI*** | 4, 44 | 13.46 | <0.01** |
| ***Cortical depth*** | 2, 22 | 4.27 | 0.06 |
| ***ROI ×<br>relative ocular polarity*** | 1, 11 | 6.67 | 0.03* |
| ***ROI ×<br>ODI*** | 4, 44 | 1.33 | 0.28 |
| ***ROI ×<br>cortical depth*** | 2, 22 | 0.54 | 0.49 |
| ***Relative ocular polarity ×<br>ODI*** | 4, 44 | 0.64 | 0.45 |
| ***Relative ocular polarity ×<br>cortical depth*** | 2, 22 | 0.65 | 0.49 |
| ***ODI ×<br>cortical depth*** | 8, 88 | 12.26 | $<10^{-4}$*** |
| ***ROI ×<br>relative ocular polarity ×<br>ODI*** | 4, 44 | 5.35 | 0.02* |
| ***ROI ×<br>relative ocular polarity ×<br>cortical depth*** | 2, 22 | 0.20 | 0.73 |
| ***ROI ×<br>ODI ×<br>cortical depth*** | 8, 88 | 3.16 | 0.04* |
| ***Relative ocular polarity ×<br>ODI ×<br>cortical depth*** | 8, 88 | 0.64 | 0.59 |
| ***ROI ×<br>relative ocular polarity ×<br>ODI ×<br>cortical depth*** | 8, 88 | 0.15 | 0.82 |

df = degrees of freedom; *: $p<0.05$; **: $p<0.01$; ***: $p<10^{-3}$

**Supplementary Table 6**. Results of applying RM-ANOVA to compare the effects of region-of-interest (ROI), relative ocular polarity, ocular dominance index (ODI), and cortical depth on interhemispheric resting-state functional connectivity (rs-FC) between central vs. peripheral V1 subregions.

| *Effect* | *df* | *F* | *p* |
|---|---|---|---|
| ***ROI*** | 1, 11 | 1.96 | 0.19 |
| ***Relative ocular polarity*** | 1, 11 | 4.46 | 0.06 |
| ***ODI*** | 4, 44 | 18.84 | $<10^{-3}$*** |
| ***Cortical depth*** | 2, 22 | 3.58 | 0.08 |
| ***ROI ×<br>relative ocular polarity*** | 1, 11 | 0.01 | 0.94 |
| ***ROI × ODI*** | 4, 44 | 3.38 | 0.08 |
| ***ROI ×<br>cortical depth*** | 2, 22 | 24.77 | $<10^{-3}$*** |
| ***Relative ocular polarity ×<br>ODI*** | 4, 44 | 1.02 | 0.35 |
| ***Relative ocular polarity ×<br>cortical depth*** | 2, 22 | 0.23 | 0.70 |
| ***ODI ×<br>cortical depth*** | 8, 88 | 20.63 | $<10^{-5}$*** |
| ***ROI ×<br>relative ocular polarity ×<br>ODI*** | 4, 44 | 0.19 | 0.70 |
| ***ROI ×<br>relative ocular polarity ×<br>cortical depth*** | 2, 22 | 4.74 | 0.02* |
| ***ROI ×<br>ODI ×<br>cortical depth*** | 8, 88 | 0.67 | 0.52 |
| ***Relative ocular polarity ×<br>ODI ×<br>cortical depth*** | 8, 88 | 0.23 | 0.82 |
| ***ROI ×<br>relative ocular polarity ×<br>ODI ×<br>cortical depth*** | 8, 88 | 3.03 | 0.048* |

df = degrees of freedom; *: $p<0.05$; ***: $p<10^{-3}$

### **Supplementary Analysis**

To better clarify the relationship between resting-state functional connectivity (rs-FC) and ocular dominance index (ODI), we fitted an linear mixed effects (LME) model to the rs-FC data using the product of the ODI of the seed and target vertices, relative ocular polarity, and cortical depth as independent factors (see Section 2.5.). In this test, rather than using the five cumulative ODI categories, all 55 possible combinations of ODIs for the seed and target vertices were used in the LME model. Consistent with the results of the RM-ANOVA, the LME results (Table S2) indicated significant effects of ODI ($t=16.83$, $p<10^{-61}$), and relative ocular polarity ($t=2.00$, $p<0.045$), as well as significant interactions between the effect of ODI and relative ocular polarity ($t=2.67$, $p<0.01$) and between the effect of ODI and middle (relative to deep) cortical depth ($t=-3.62$, $p<10^{-3}$) as well as superficial (relative to deep) cortical depth ($t=-7.17$, $p<10^{-12}$).
